# FLI1 localization to the chlamydial inclusion involves multiple mechanisms

**DOI:** 10.1101/2023.10.17.562819

**Authors:** Natalie A. Sturd, Macy G. Wood, Legacy Durham, Scot P. Ouellette, Elizabeth A. Rucks

## Abstract

Following entry into a host cell, the obligate intracellular pathogen, *Chlamydia trachomatis*, establishes an intracellular niche within a membrane derived vacuole called the chlamydial inclusion. The resulting inclusion membrane is modified by the pathogen and is a hybrid host-chlamydial structure. From within this intracellular niche, *C. trachomatis* must orchestrate numerous host-pathogen interactions to surreptitiously acquire nutrients from its host and to limit detection by the host innate immune system. *C. trachomatis* mediates many of these interactions with the host, in part, by using a family of type III secreted membrane proteins, termed inclusion membrane proteins (Incs). Incs are embedded within the inclusion membrane, and some function to recruit host proteins to the inclusion. Two such recruited host proteins are leucine <u>r</u>ich <u>r</u>epeat <u>F</u>lightless-1 interacting <u>p</u>rotein 1 (LRRF1/LRRFIP1) and its binding partner Flightless 1 (FLI1/FLII). LRRF1 interacts with Inc protein Ct226. However, interactions of FLI1 with candidate Incs or with LRRF1 during infection have not been defined. We hypothesized that FLI1 recruitment to the inclusion would be dependent on LRRF1 localization. To test this hypothesis, we used siRNA targeting *lrrf1* or *fli1,* revealing that FLI1 can localize to the inclusion independently of LRRF1. Therefore, to further characterize FLI1 localization, we developed and characterized a series of CRISPRi knockdown and complementation strains in *C. trachomatis* serovar L2 that target *ct226* and co-transcribed candidate Incs, *ct225* and *ct224*, to understand the mechanisms of FLI1 and LRRF1 localization to the inclusion. Our results indicate that FLI1 is recruited to the inclusion by multiple mechanisms.

**IMPORTANCE:** *Chlamydia trachomatis* is a leading cause of both preventable infectious blindness and bacterial sexually transmitted infections worldwide. Since *C. trachomatis* must grow and replicate within human host cells, it has evolved several ways of manipulating the host to establish a successful infection. As such, it is important to describe the interactions between host proteins and chlamydial proteins to understand which strategies *C. trachomatis* uses to shape its intracellular environment. This study looks in detail at such interactions of two host proteins, FLI1 and LRRF1, during chlamydial infection. Importantly, the series of knockdown and complement strains developed in this study suggest these proteins have both independent and overlapping mechanisms for localization, which ultimately will dictate how these proteins function during chlamydial infection.

## INTRODUCTION

In the year 2020, the World Health Organization reported 129 million new *Chlamydia trachomatis* infections worldwide with global infection rates rising steadily since 2013 (1). In the United States alone, direct medical costs of treating infections exceeds 650 million dollars, second only to direct medical costs of HPV and HIV (2). Clinical manifestations of *C. trachomatis* infections are serovar-dependent, as serovars dictate tissue tropism. *C. trachomatis* is capable of infecting multiple host tissues, including the conjunctiva of the eye (serovars A-C), the male and female reproductive tracts (serovars D-K), and the pelvic lymph nodes, (serovars L1-L3), with the L serovars causing lymphogranuloma venereum (3, 4). Importantly, studies estimate upwards of 70-80% of infected women and 50-60% of infected men are asymptomatic (5, 6), which often results in a delay of treatment and resolution of infection. Chronic infection in the female reproductive tract increases the risk of the infection ascending into the uterus and fallopian tubes, leading to pelvic inflammatory disease (PID)(7). Chronic and repeat chlamydial infections increase the risk of uterine and tubal scarring which have the potential to progress to tubal factor infertility (TFI) or hospitalization for ectopic pregnancy (7–10). The host-pathogen interactions responsible for the development of host tissue pathology are not fully understood.

*C. trachomatis* is a gram-negative, obligate intracellular pathogen, and requires infection of eukaryotic cells to complete its developmental cycle. It alternates between two morphologically and functionally distinct forms: the infectious, non-replicative elementary body (EB) and the non-infectious, replicative reticulate body (RB)(11, 12). The developmental cycle begins with entry of the EB via host-mediated endocytosis, where it resides in a host-membrane derived vacuole. Following entry, the EB rapidly undergoes primary differentiation to an RB, which initiates replication within this pathogen-specific vacuole, termed the chlamydial inclusion (13). Later in development, RBs undergo secondary differentiation to EBs (12), which egress either by host cell lysis or by extrusion to infect new host cells (14). Gene expression throughout the developmental cycle is temporally regulated, and gene transcription profiles are categorized by peak expression into early, mid, and late-cycle development (15, 16). Throughout the developmental cycle, *C. trachomatis* must maintain and grow the inclusion, acquire nutrients, and limit detection by the host until it escapes from the cell. All this requires complex and tightly regulated interactions with host proteins and associated signaling pathways while remaining sequestered within the inclusion.

*C. trachomatis* uses a type three secretion system to secrete effectors to establish and maintain their intracellular niche (17, 18). One such family of secreted effectors is known as inclusion membrane proteins, or Incs, which localize within the inclusion membrane (IM) (19). Incs are characterized as having at least two transmembrane domains with the N-and C-termini of the protein oriented towards the host cell cytoplasm (20). While some Incs likely support the structural integrity and shape of the inclusion membrane, most studies characterizing Inc function have focused on interactions between Incs and various host proteins (21–25). These Inc-host interactions allow for selective interaction with host cell vesicular compartments, acquisition of host-derived lipids, and avoidance of innate immune defenses (26, 27). However, the full scope of Inc-host and/or Inc-Inc interactions and their impact on host pathways remains unknown.

Previously, we used APEX2 proximity labeling to determine the host protein interactome around the inclusion (28, 29). Two statistically significant hits were the eukaryotic protein leucine <u>r</u>ich <u>r</u>epeat <u>F</u>lightless-1 interacting <u>p</u>rotein 1 (**LRRF1/**LRRFIP1/GCF2) and its binding partner, Flightless-1 (**FLI1**/FLII), both of which localize to the inclusion membrane during chlamydial infection (28). Further, LRRF1 localizes to the inclusion during early mid-cycle development (∼12 hours post infection (hpi)) and interacts with the chlamydial Inc protein Ct226 (28). FLI1 and LRRF1 have a multitude of reported functions in uninfected eukaryotic cells. Both are reported to act as regulators of eukaryotic protein expression; FLI1 is a co-activator of nuclear receptor mediated transcription (e.g., estrogen receptor and glucocorticoid receptor) (30–32), while LRRF1 is a known transcriptional repressor with DNA binding activity (e.g., *tnf*) (33–36). Both are also involved in regulating host innate immune responses to infection, though their activity has been shown to be both antagonistic and synergistic (37–43). Finally, both proteins are involved in modulation of the actin cytoskeleton. FLI1 binds and prevents actin polymerization and is a prominent negative regulator of fibroblast migration and wound healing (44–50). In contrast, knockdown of LRRF1 reduces migration, cytoskeletal remodeling, and RhoA activation, potentially through its ability to interact with the RhoA-activator LARG (45, 51). Considering the respective activities of FLI1 and LRRF1, it is intriguing that they localize to the inclusion membrane during *C. trachomatis* infection. However, interactions of FLI1 with candidate Incs or with LRRF1 during infection have not been clearly defined. Furthermore, our understanding of FLI1 and LRRF1 function during chlamydial pathogenesis and of their overall contribution to host pathology has been difficult to characterize.

Therefore, in this study, we investigated the dynamics of FLI1 localization to the chlamydial inclusion membrane throughout the developmental cycle and interrogate the Inc-host interactions required for both FLI1 and LRRF1 localization. Our data demonstrate that FLI1 interacts with Ct226 in a complex only in the presence of LRRF1 but cannot bind Ct226 independently. Further, FLI1 can localize to the chlamydial inclusion independently of LRRF1, indicating a LRRF1-/Ct226-independent mechanism of FLI1 localization. To better understand FLI1 and LRRF1 localization dynamics, we used a CRISPR inference (CRISPRi) inducible knockdown system in *C. trachomatis* serovar L2 (Ctr L2) (52, 53) and developed a cadre of knockdown and complement strains targeting *ct226* and co-transcribed genes *ct225* and *ct224*. Lastly, our data strongly suggest that Ct225 is not an Inc, as previously annotated in the literature, and instead localizes within the bacterial membrane during infection. Ultimately our data demonstrate that there are multiple mechanisms for FLI1 recruitment, which may have implications for FLI1 function at the IM.

## RESULTS

### Characteristics of FLI1 localization to the inclusion

Our previous study determined that LRRF1 recruitment to the inclusion membrane begins during mid-cycle (by 12hpi) and remains at the inclusion through late cycle timepoints (examined through 36hpi) (28). We hypothesized that FLI1 follows a similar timeline for localization. To determine when FLI1 localizes to the inclusion membrane, HEp2 cells were infected with wild-type *C. trachomatis* serovar L2 (Ctr L2), then fixed at time points between 8 and 46hpi and processed for indirect immunofluorescence (IFA). FLI1 first localized to the inclusion between 12-14 hpi and remained stably associated with the inclusion throughout the developmental cycle, up to 46hpi (Figure 1A).

**Figure 1.**
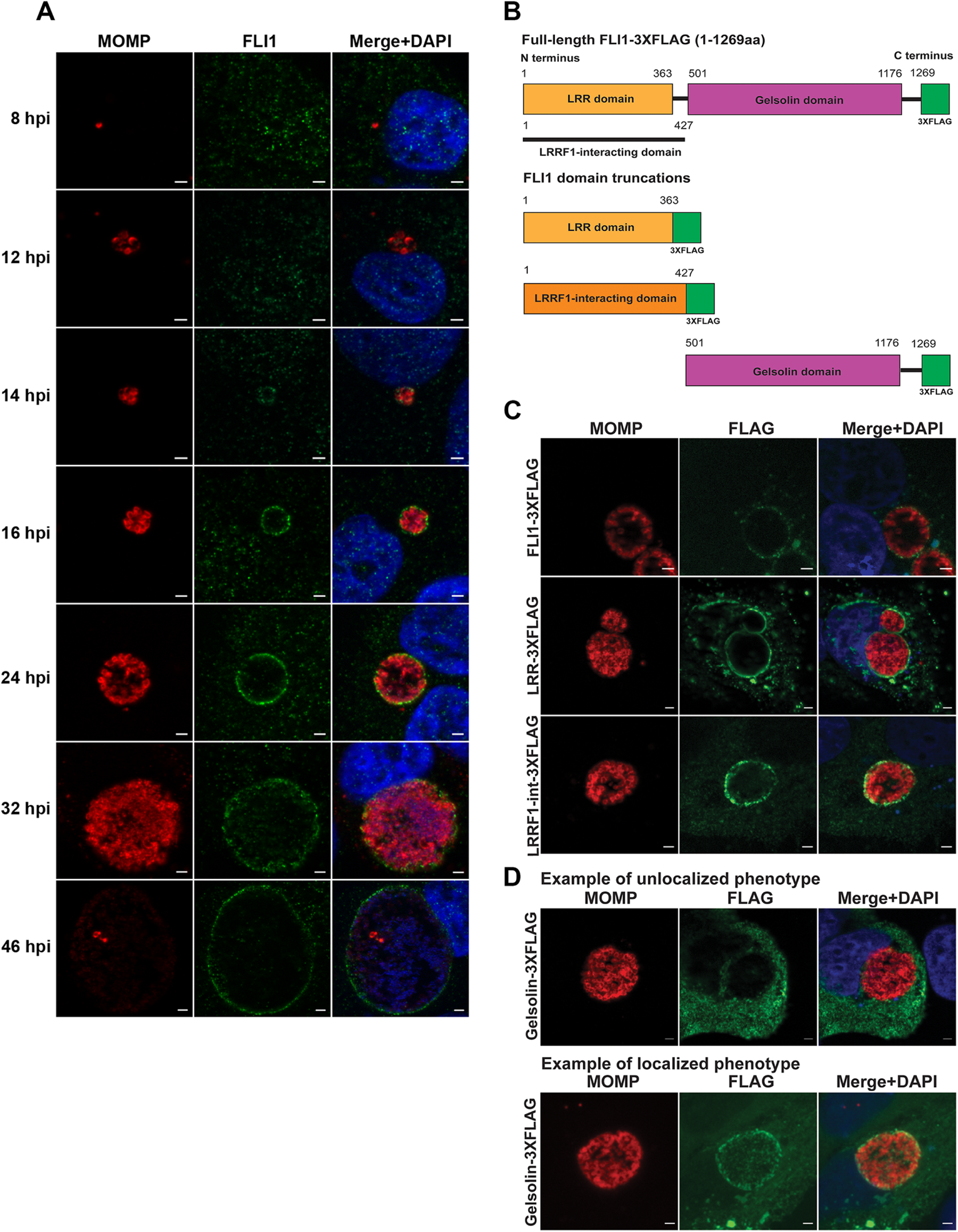
Characteristics of FLI1 localization to the chlamydial inclusion. (A) Time-course of Flightless 1 (FLI1) recruitment to the inclusion. HEp2 cells were plated and infected with wild-type *Chlamydia trachomatis* serovar L2, as described in Methods and Materials. Coverslips were recovered and fixed with 4% paraformaldehyde at the indicated timepoints. Fixed coverslips were permeabilized with methanol and stained for indirect immunofluorescence to visualize FLI1 (anti-FLI1; green), chlamydial organisms (anti-MOMP; red), and DNA (DAPI; blue). FLI1 is recruited to the inclusion membrane as early as 12 hpi with more robust localization apparent at 14 hpi. FLI1 localization at the IM remains stable throughout infection up to 46 pi. Scale bar=2µm. (B) Diagram of domain truncations of FLI1. Full-length FLI1 and truncation constructs were created to determine which domains of FLI1 are required for inclusion localization. Full length FLI1-3XFLAG (residues 1-1269aa) contains the LRR domain and the gelsolin domain. Residues 1-427aa are necessary for interaction with LRRF1. Domain truncations of FLI1. Constructs LRR-3XFLAG (residues 1-363) and LRRF-int-3XFLAG (residues 1-427aa) contain the LRR domain but truncates the gelsolin domain. The Gelsolin-3xFLAG (507-1269aa) construct truncates the LRR domain. (C-D) Localization of truncation mutants during Ctr L2 infection. HEp2 cells were plated, transfected with the indicated construct and infected with wild-type Ctr L2. Cells were fixed at 24hpi and processed for immunofluorescence to visualize chlamydial organisms (anti-MOMP; red), full-length or truncated FLI1 (anti-FLAG; green), and DNA (DAPI; blue). (C) Full length FLI1-3XFLAG, LRR-3xFLAG, and LRRF1-int-3XFLAG localized to wild-type inclusions. (D) The majority of inclusions lacked Gelsolin-3XFLAG labeling and instead remained in the cell cytoplasm (unlocalized phenotype), but a small percentage of inclusions demonstrated a localized phenotype.

FLI1 contains an N-terminal leucine rich repeat (LRR) domain and a C-terminal gelsolin domain that are responsible for protein interactions and actin cytoskeleton binding/remodeling, respectively (54, 55). It can also interact with LRRF1 via its LRR domain (56, 57). To assess which domains are required for localization to the inclusion, we created a full-length FLI1-3XFLAG construct (Figure 1B) alongside three truncation constructs: the LRR-3XFLAG (1-363aa), Gelsolin-3XFLAG (501-1269aa), and a truncation construct containing the region of FLI1 required for LRRF1 interaction, LRRF1-int-3XFLAG (1-427aa), which includes the LRR domain (Figure 1B). The constructs were transfected into HEp2 cells, which were subsequently infected with wild-type Ctr L2. Cells were fixed and processed at 24hpi for immunofluorescence microscopy. Full-length FLI1-3XFLAG, LRR-3XFLAG, and LRRF1-int-3XFLAG were all shown to localize to the inclusion membrane (Figure 1C). In contrast, most inclusions lacked Gelsolin-3XFLAG labeling, though a small percentage of inclusions demonstrated a localized phenotype (Figure 1D). This suggests that while the LRR and LRRF1-interacting domains of FLI1 reliably localize to the inclusion membrane, there may be an alternative, albeit less efficient, mechanism for FLI1 recruitment via its gelsolin domain.

### FLI1 and LRRF1 localize independently of each other yet interact together in complex with Ct226

LRRF1 and FLI1 are known to interact via their respective leucine-rich repeat (LRR) domains (55, 57, 58) and both proteins localize to the inclusion around the same point in development (between 12 and 14hpi). Additionally, LRRF1 interacts with chlamydial Inc protein Ct226 and overexpression of Ct226-FLAG increases both LRRF1 and FLI1 recruitment to the inclusion (28). Therefore, we hypothesized that localization of both proteins may be dependent on each other. To test this, we performed siRNA knockdown of either FLI1 or LRRF1, infected with wild-type Ctr L2, and used IFA to determine localization of the other protein. Following siRNA knockdown of LRRF1, FLI1 localized to the inclusion membrane (Figure 2A; Supplemental Figure 1A). Similarly, LRRF1 localized to the inclusion in the absence of FLI1 (Figure 2B; Supplemental Figure 1B). These data suggest that FLI1 and LRRF1 can localize to the inclusion membrane independently of each other.

**Figure 2.**
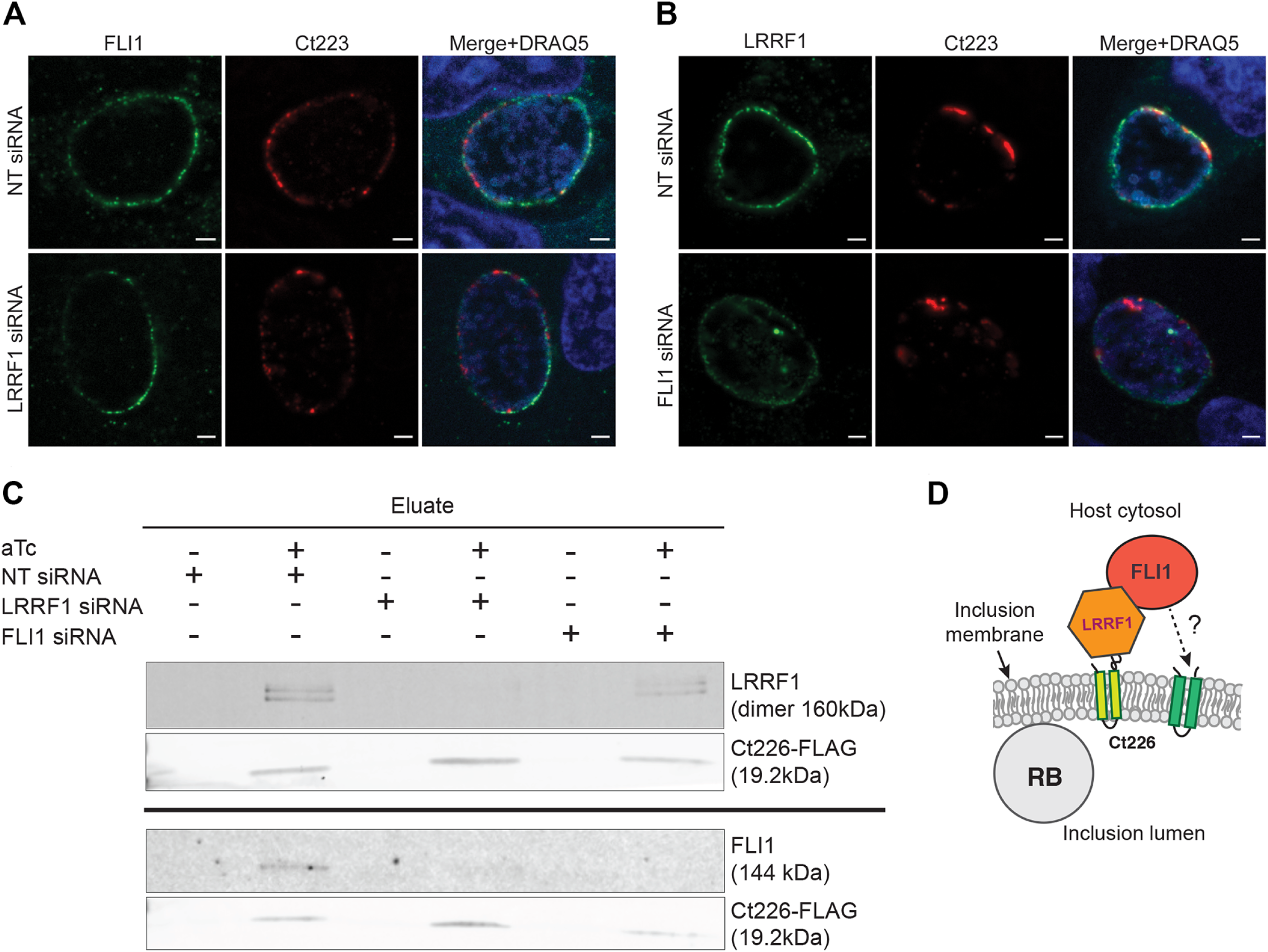
FLI1 and LRRF1 localize independently of each other to the inclusion membrane but interact in complex with Inc protein Ct226. (A-B) HeLa cells were treated with either non-targeting NT, LRRF1, or FLI1 siRNA, infected with wild-type Ctr serovar L2 and fixed with 4% paraformaldehyde at 24hpi. Indirect immunofluorescence was used to visualize FLI1 (anti-FLI1; green) or LRRF1 (anti-LRRF1; green), and Inc protein Ct223 (anti-Ct223; red), which is organized in microdomains around the inclusion. DRAQ5 was used to visualize host and bacterial DNA. (A) FLI1 localizes to the inclusion membrane following both treatment with NT and LRRF1 siRNA. (B) LRRF1 localizes to the inclusion membrane following both treatment with NT and FLI1 siRNA. Scale bar=2µm. (C) Co-immunoprecipitation of FLI1 and LRRF1 with Ct226-FLAG. HEp2 cells were treated with either non-targeting (NT) siRNA, FLI1 siRNA, or LRRF1 siRNA followed by infection with *C. trachomatis* serovar L2 carrying pBOMB4-Ct226-FLAG. Expression of Ct226-FLAG was induced with 5nM aTc at 7hpi and cell lysates were collected at 24hpi for co-immunoprecipitation as described in Methods and Materials. Eluate fractions were blotted for FLI1 (144 kDa), LRRF1 (dimer, 160 kDa), and Ct226-FLAG (19.2 kDa). Both LRRF1 and FLI1 are detected in the eluate fractions of cells treated with NT siRNA. However, FLI1 does not appear in the eluate fraction of cells treated with LRRF1 siRNA. LRRF1 is detected of the eluate fraction when cells are treated with FLI1 siRNA. These data are consistent with LRRF1 interacting directly with Ct226, while FLI1 cannot interact in complex with Ct226 independent of LRRF1. (D) Diagram to illustrate the FLI1-LRRF1-Ct226 interactions at the inclusion.

Next, we tested whether FLI1 can interact with Ct226 by infecting HEp2 cells with a Ctr L2 strain carrying a plasmid with an inducible Ct226-FLAG construct, pBOMB4-*ct226-FLAG*. Ct226-FLAG expression was induced or not at 7hpi, and Ct226-FLAG was affinity purified from lysates collected at 24hpi. FLI1 was detected in the eluate fraction of induced samples, indicating that it can interact either directly or indirectly with Ct226 (Supplemental Figure 2). To determine if FLI1 requires LRRF1 to interact with Ct226, HEp2 cells were treated with siRNA targeting LRRF1 or FLI1 then infected with the strain carrying pBOMB4-*ct226-FLAG*. Cell lysates were collected at 24hpi, and Ct226-FLAG was affinity purified. LRRF1 co-immunoprecipitated with Ct226 following FLI1 knockdown (Figure 2C). However, FLI1 did not co-immunoprecipitate with Ct226 following LRRF1 knockdown (Figure 2C). These results indicate FLI1 cannot interact with Ct226 independently of LRRF1 (Figure 2C & 2D) even though it is recruited to the inclusion membrane in the absence of LRRF1 (Figure 2A). These data suggest that there may be additional Incs that interact with FLI1 and mediate its localization.

### Chlamydial knockdown and complement strains targeting Ct226 expression

To test the hypothesis that knockdown of *ct226* would inhibit recruitment of LRRF1, but not FLI1, to the chlamydial inclusion, we created an inducible knockdown strain targeting *ct226* using a dCas12-based CRISPR interference (CRISPRi) inducible knockdown system (53). The principle of CRISPRi relies on a catalytically dead Cas12 enzyme, which is guided to the target genetic sequence by a CRISPR RNA (crRNA) (52, 53) to sterically inhibit transcription at that site. Into the pBOMBL12CRia vector, we inserted a crRNA targeting the intergenic region upstream of *ct226* to create the vector pBOMBL12CRia(*ct226*) (Figure 3A) and transformed the resulting plasmid into Ctr L2, herein referred to as the L2/*ct226* KD strain. We also included the strain carrying the empty vector plasmid, pBOMBL12CRia(*empty vector*) (*E.V.*), as the negative control in these studies, which maintains inducible expression of dCas12 but lacks a targeting crRNA. To characterize the strains, we validated knockdown using RT-qPCR to measure transcript levels and performed inclusion forming unit (IFU) assays to determine if chlamydial development was grossly affected.

**Figure 3.**
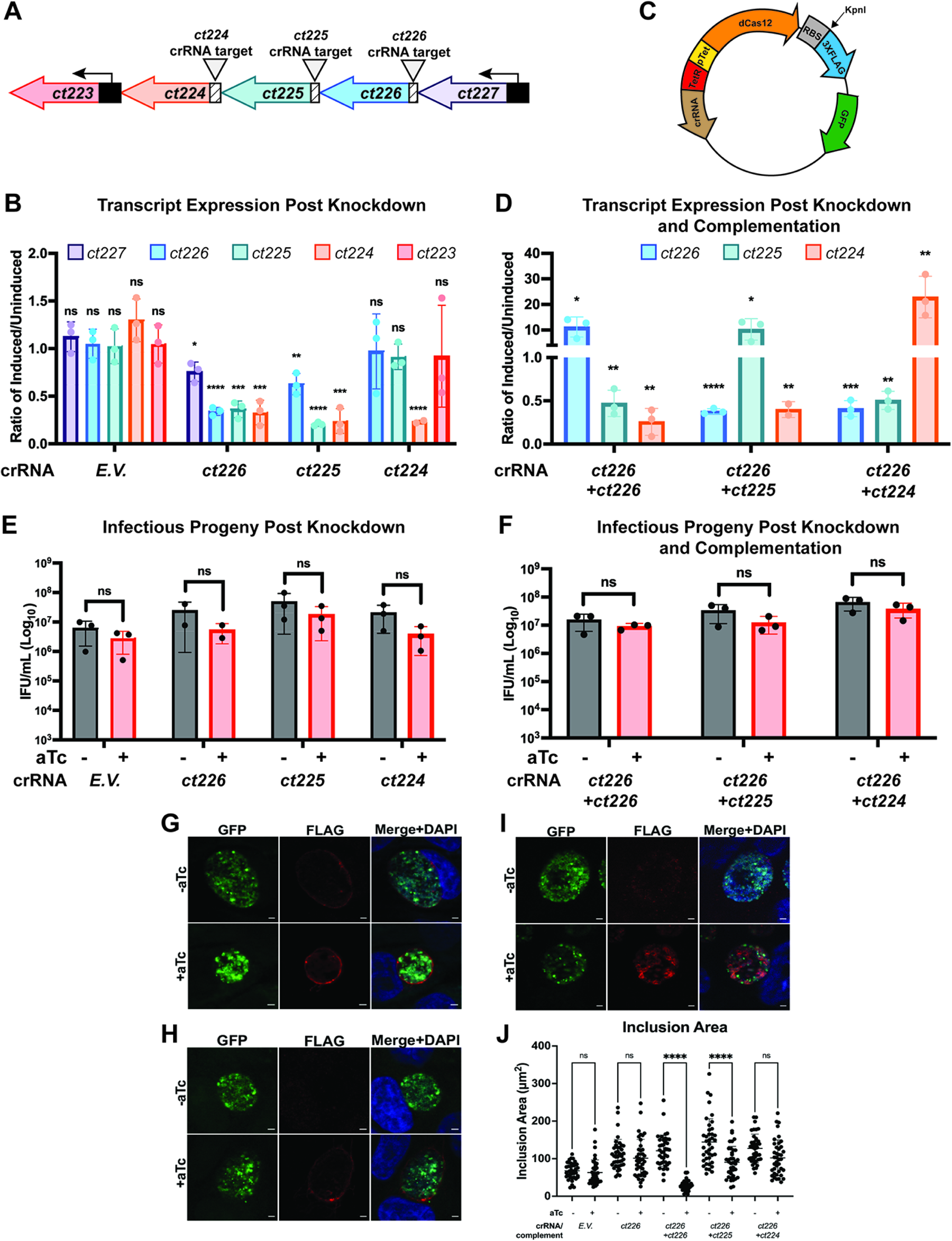
Characterization of knockdown and complementation strains by RT-qPCR, infectious progeny production, and indirect immunofluorescence microscopy. (A) Diagram of the *ct227* operon. Triangles indicate the binding site of the indicated CRISPR RNA (crRNA) in the intergenic region upstream of the targeted gene. (B) RT-qPCR analysis of knockdown strains. HEp2 cells were seeded, infected with the indicated strains, and induced as described in Materials and Methods. RNA was collected and processed for RT-qPCR. cDNA transcript levels of genes were normalized to 16s transcripts and reported as the ratio of uninduced to induced at 12hpi. A paired student’s t-test was used to determine statistically significant differences in transcript levels between uninduced and induced conditions for each strain. (C) Diagram of vector design for complementation downstream of dCas12 in pBOMBL12CRia vector. (D) RT-qPCR analysis of complement strains. Samples were collected and analyzed as described above. (E) Infectious progeny production in knockdown strains. Infectious progeny were measured in both uninduced and induced inclusions and reported as inclusion forming units (IFU) per mL and a paired student’s t-test was used to determine statistical significance between uninduced and induced conditions for each strain. (F) Infectious progeny production in complementation strains. (G-I) Indirect immunofluorescence confirming expression and localization of the complemented 3XFLAG-tagged Inc for all three complement strains. HEp2 cells were seeded on glass coverslips, infected with appropriate complement strain and induced as described in the Materials and Methods. At 24hpi, the coverslips were processed for indirect immunofluorescence to visualize the 3XFLAG-tagged Inc (anti-FLAG; red) and host and chlamydial DNA (DAPI; blue). Constitutively expressed GFP was used to visualize the chlamydial organisms (GFP; green). (G) Ct226-3XFLAG is dimly visible at the inclusion under un-inducing conditions, indicating some leaky expression of the complement. Under induced conditions, Ct226-3XFLAG was secreted and localized as expected to the chlamydial inclusion membrane. (H) Complemented Ct224-3XFLAG is secreted and localizes to the inclusion membrane, as anticipated. (I) Ct225-3XFLAG notably localizes to the chlamydial organisms, as opposed to the inclusion membrane. (J) Quantification of inclusion area in complementation strains by ImageJ. An ordinary two-way ANOVA test was performed to identify statistically significant differences in inclusion area between the complement strains compared to the negative control strain and the L2/*ct226* KD strain. Neither E.V. strain nor L2/*ct226* KD strain demonstrated a difference in inclusion size between un-inducing and inducing conditions. Complementation of either Ct226 or Ct225 led to a 77% or 37% decrease in inclusion area, respectively, while complementation of Ct224 did not impact inclusion area. Scale bar= 2μm. Statistical significance was reported as follows: *p<0.05, **p<0.01, ***p<0.001, ****p<0.0001, ns= not significant.

To measure transcript levels and validate knockdown, HEp2 cells were infected with either L2/*ct226* KD or the E.V. control strain, and knockdown was induced using anhydrotetracycline (aTc) as described in Materials and Methods. RNA was collected at 3 (time of aTc addition), 12 (peak transcription of *ct226*), and 24 (*ct226* transcription returns to baseline) hpi, which also captures transcription at early, mid, and late-cycle chlamydial development, respectively (15). *ct226* is located in the *ct227* operon, where genes *ct227, ct226, ct225,* and c*t224* are co-transcribed (Figure 3A). Therefore, genes upstream and downstream of *ct226* were also measured to determine any polar effects of knockdown. Expression of dCas12 alone did not result in gene knockdown as the negative control strain did not demonstrate any differences in transcript levels of the *ct227* operon (Figure 3B) following induction. For the L2/*ct226* KD strain, RT-qPCR analysis demonstrated ∼66% knockdown of *ct226* transcripts at 12hpi when compared with the uninduced samples (Figure 3B). Transcripts for downstream genes *ct225* and *ct224* also showed 64% and 67% knockdown, respectively, and transcripts of upstream gene *ct227* demonstrated a 24% decrease compared to the uninduced levels (Figure 3B). Therefore, targeting *ct226* by CRISPRi significantly represses transcription of *ct226*, *ct225*, and *ct224*.

Given the polar effects on *ct225* and *ct224*, we created and characterized two additional knockdown strains specifically targeting these genes (Figure 3A), herein referred to as L2/*ct225* KD and L2/*ct224* KD strains, respectively. Following induction of L2/*ct225* KD strain, both *ct225* (∼79%) and *ct224* (∼76%) transcripts were decreased (Figure 3B) at 12hpi. Additionally, a 37% decrease in *ct226* transcripts was observed, but Ct226 protein levels were unchanged by western blot analysis (Figure 3B; Supplemental Figure 3). Induction of the L2/*ct224* KD strain resulted in a 77% decrease in *ct224* transcripts while *ct226* and *ct225* transcript levels were unchanged. Of note, targeting *ct224* with CRISPRi does not alter transcript levels of *ct223* (Figure 3B), which resides in the operon immediately downstream.

Additionally, to assess the contribution of each Inc in the absence of the others, we individually complemented *ct226, ct225,* or *ct224* in the L2/*ct226* KD background. For complementation, we modified the pBOMBL12cRia(*ct226*) vector by inserting a KpnI digest site and 3XFLAG tag directly downstream of dCas12 (Figure 3C). The KpnI site allowed complementation of each individual Inc, under the same inducible promoter as dCas12. By RT-qPCR analysis, induction of dCas12 and complementation of Ct226-3XFLAG in the L2/*ct226* KD strain background led to an ∼8.7-fold increase in *ct226* transcripts and an 62.9% and 81.6% decrease in *ct225* and *ct224* transcripts, respectively, compared to the uninduced samples (Figure 3D). Complementation of Ct225-3XFLAG led to a 10-fold increase in *ct225* transcripts compared to uninduced conditions whereas both *ct226* (62% decrease) and *ct224* (60% decrease) transcript levels decreased in inducing conditions (Figure 3D). Finally, complementation of Ct224-3XFLAG led to a ∼23 fold increase in *ct224* transcripts following induction, while *ct226* and *ct225* transcripts were decreased 59% and 49%, respectively (Figure 3D).

To determine if CRISPRi targeting of these genes would grossly impact chlamydial development, we performed inclusion forming unit (IFU) assays. The principle of IFU assays for *C. trachomatis* is analogous to the colony forming unit (CFU) assay, as it assesses the number of viable EBs produced from a primary infection. HEp2 cells were infected with the indicated strains and induced as described in Materials and Methods. EBs from the primary infection were collected and used to infect fresh monolayers, and inclusions from this secondary infection were quantified at 24hpi (Figures 3E & 3F). Notably, induction of knockdown in the E.V. control strain demonstrated a ∼48% decrease in infectious progeny (uninduced 6.08x10^6^ IFU/mL; induced 2.79x10^6^ IFU/mL; Figure 3E), consistent with increased energy requirements likely associated with dCas12 expression (59). For the L2/*ct226* KD strain, infectious progeny were reduced by 69% following knockdown (uninduced 1.87x10^7^ IFU/mL; induced 4.14x10^6^ IFU/mL; Figure 3E). This represents a mild decrease in infectious progeny compared to the negative control and is not considered biologically significant (i.e., 1-log reduction in IFUs). Induction of the L2/*ct225* KD strain resulted in a 60% reduction in IFUs (uninduced 4.82x10^7^ IFU/ml; induced 1.77x10^7^ IFU/mL) when compared to uninduced conditions (Figure 3E). Knockdown of *ct224* resulted in an 80% decrease in IFUs (uninduced 2.04x10^7^ IFU/ml; induced 3.80x10^6^ IFU/ml; Figure 3E), which is the greatest reduction across all three knockdown strains but is still not considered biologically relevant. Induction of knockdown and complementation in the strain carrying pBOMBL12cRia(*ct226*)-*ct226-3XFLAG* resulted in a 48.6% decrease in the number of infectious progeny (uninduced 1.54x10^7^ IFU/ml; induced 8.97x10^6^ IFU/ml; Figure 3F). Recovered infectious progeny following complementation of Ct225-3XFLAG demonstrated a 46% decrease (uninduced 3.28x10^7^ IFU/mL; induced 1.20x10^7^ IFU/mL; Figure 3F). Induction of complementation of Ct224-3XFLAG yielded a 58% decrease in infectious progeny compared to uninduced inclusions (4.81x10^7^ IFU/mL; induced 2.74x10^7^ IFU/mL; Figure 3F). Overall, these data suggest that knocking down any or all of *ct226, ct225*, or *ct224* has a minimal impact on developmental cycle progression.

Lastly, we used IFA to assess expression and localization of the complemented Inc proteins. Complementation of Ct226-3XFLAG demonstrated some leaky expression under non-inducing conditions both by microscopy and by western blot (Figure 3G; Supplemental Figure 4). Following induction, Ct226-3XFLAG is expressed and secreted to the inclusion membrane (Figure 3G). Ct224-3XFLAG was also secreted and inserted into the inclusion membrane and appeared to localize in microdomains (a pattern of localization where Incs localize in discrete clusters within the inclusion membrane (60)) in the inclusion membrane (Figure 3H). Surprisingly, IFA revealed that Ct225-3XFLAG did not localize to the inclusion membrane and instead localized to the bacterial membrane (Figure 3I). This was unexpected as Ct225 has been identified in the literature as an Inc protein (61, 62). To ensure that overexpression of Ct225 in the absence of Ct226 or Ct224 was not preventing type III secretion of Ct225, we created an inducible overexpression strain encoding *ct225-FLAG.* We performed induction tests of Ct225-FLAG expression using low aTc concentrations (0.5nM aTc) and found that Ct225-FLAG still localized to the bacterial membrane (Supplemental Figure 5A & 5B). For additional rigor, we performed immunofluorescence assays using an anti-Ct225 antibody to assess endogenous Ct225 localization in both wild-type *C. trachomatis* serovar L2 (Ctr L2) and the Ct225-FLAG overexpression strain. In Ctr L2, endogenous Ct225 localized to the organism membrane, and it co-localized with the organism-associated FLAG staining in the Ct225-FLAG overexpression strain (Supplemental Figure 5C and 5D). Taken together, these data suggest that Ct225 is not an Inc protein and may serve a different function in the chlamydial bacterial membrane.

**Figure 4.**
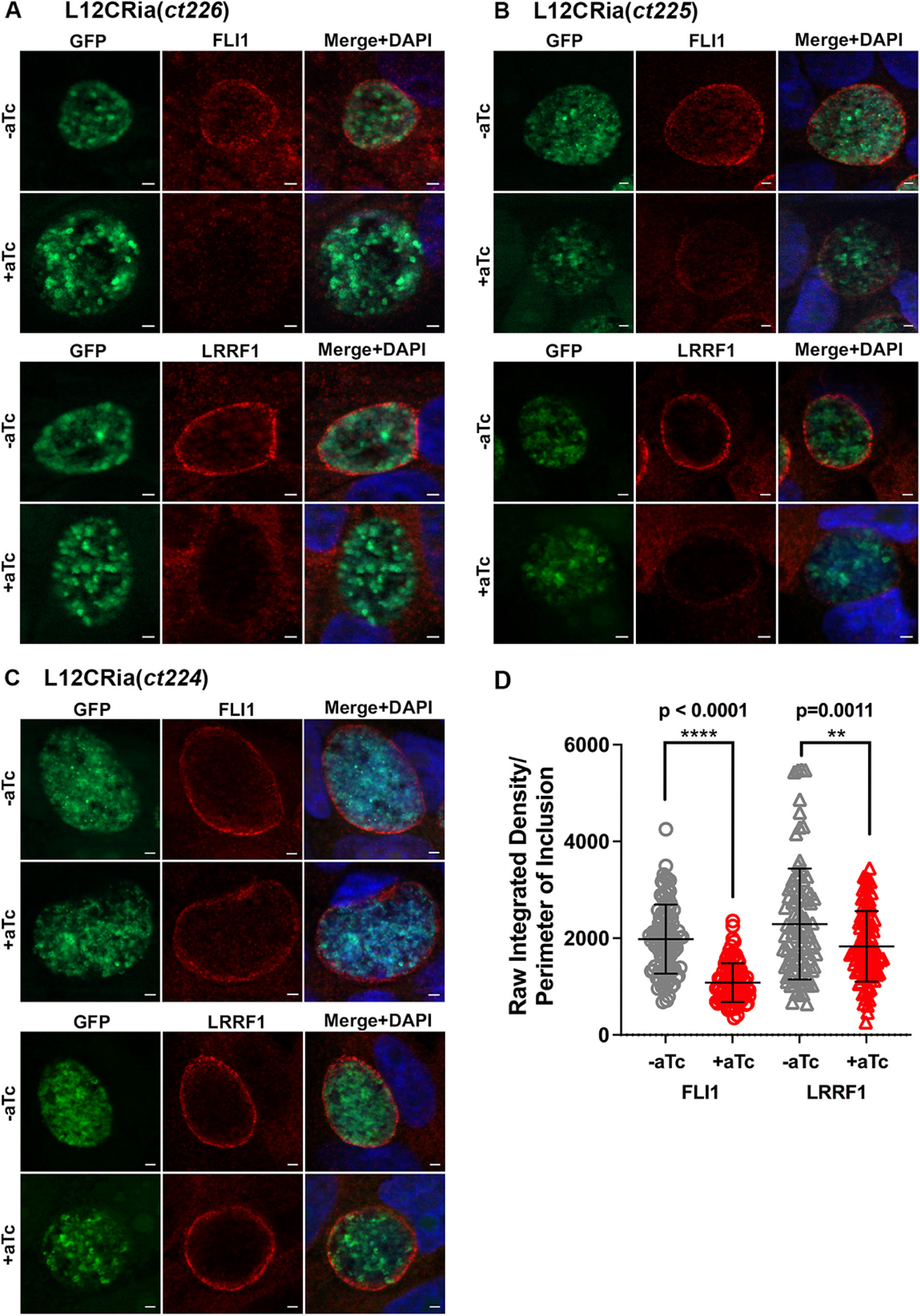
Knockdown of *ct226* and *ct225*, but not *ct224*, negatively impact localization of FLI1 and LRRF1 to the chlamydial inclusion membrane. (A-C) Localization of FLI1 and LRRF1 in knockdown strains. HEp2 cells were seeded onto glass coverslips, infected with one of the three knockdown strains or the empty vector control strain, and induced as described in the Materials and Methods. At 24hpi, coverslips were fixed and processed for indirect immunofluorescence to visualize chlamydial organisms (GFP; green), FLI1 (anti-FLI1; red) or LRRF1 (anti-LRRF1; red), and host and bacterial DNA (DAPI; blue). (A) For the L2/*ct226* KD strain, uninduced inclusions demonstrated localization of both FLI1 and LRRF1. Following knockdown, neither FLI1 nor LRRF1 localize to the inclusion membrane during infection. (B) For the L2/*ct225* KD strain, uninduced conditions again demonstrated localization of both FLI1 and LRRF1. Following knockdown, FLI1 and LRRF1 do localize to the inclusion, but fluorescence intensity of both proteins was markedly decreased compared to uninduced inclusions. (C) For the L2/*ct224* KD strain, inclusions in both the uninduced and induced conditions demonstrated FLI1 and LRRF1 localization. Scale=2um. (D) ImageJ quantification of fluorescence intensities for FLI1 and LRRF1 for the L2/*ct225* KD strain. Results were normalized to inclusion perimeter and an unpaired student’s t-test was performed to determine statistically significant differences for each protein. Knockdown led to a significant decrease in FLI1 fluorescent intensity at the inclusion membrane (****p-value<0.0001). LRRF1 also demonstrated a statistically significant decrease (**p-value=0.0011). Overall, these results suggest that induction of knockdown in the strains targeting *ct226* and *ct225* negatively impacts localization of FLI1 and LRRF1.Statistical significance is indicated as follows: *p<0.05, **p<0.01, ***p<0.001, ****p<0.0001, ns= not significant. Scale bar=2μm.

**Figure 5.**
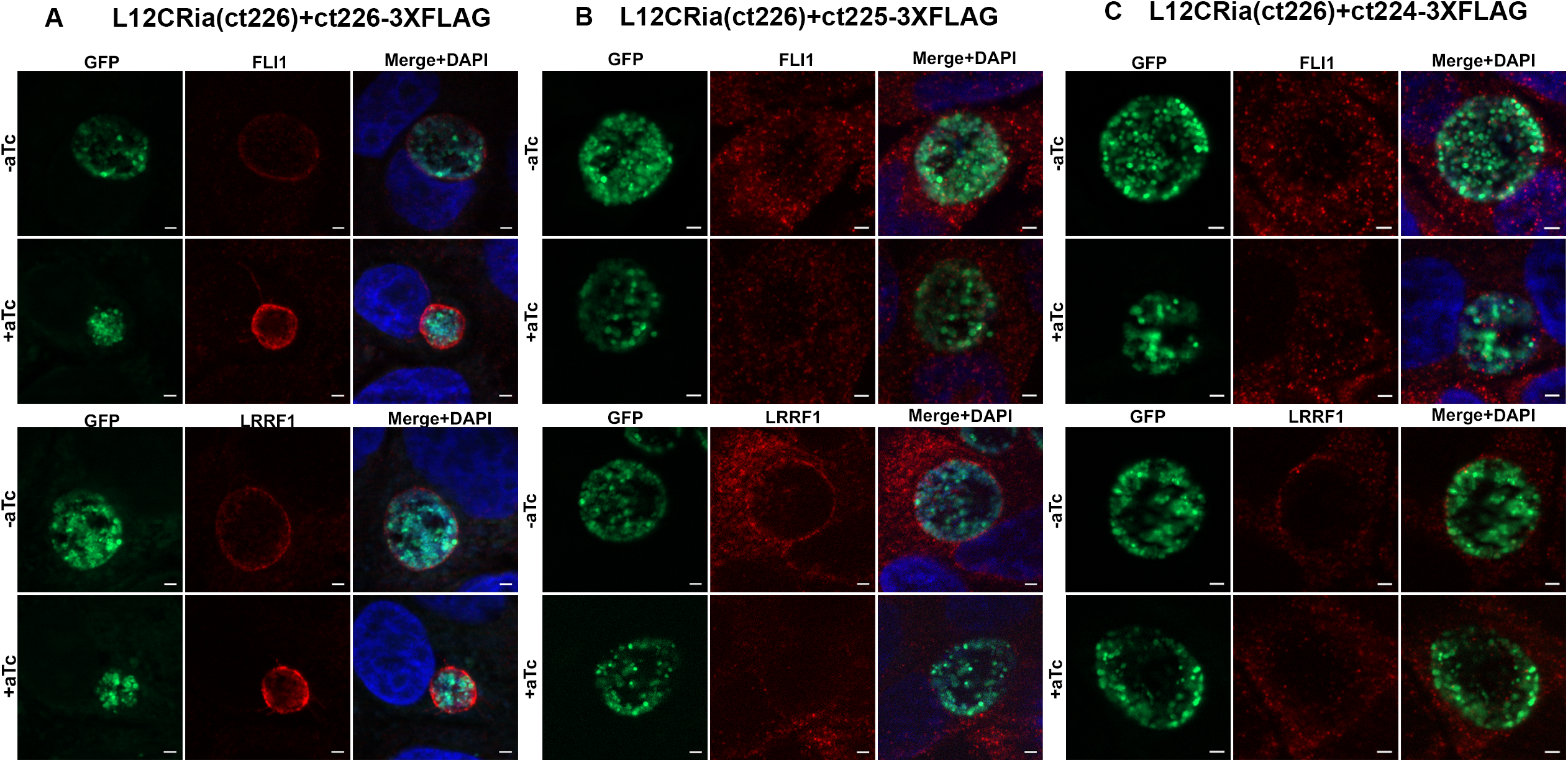
Complementation of Ct226, but not Ct225 nor Ct224, fully restores both LRRF1 and FLI1 localization to the chlamydial inclusion. (A-C) Localization of FLI1 and LRRF1 in complementation strains. HEp2 cells were seeded onto coverslips and were infected with one of the three complement strains and induced or not according to the Materials and Methods. At 24hpi, coverslips were fixed and processed for indirect immunofluorescence to visualize the chlamydial organisms (GFP; green), FLI1 (anti-FLI1; red) or LRRF1 (anti-LRRF1; red), and host and bacterial DNA (DAPI; blue). Images were taken using a Zeiss ApoTome.2 fluorescence microscope at 100x magnification. (A) For the strain carrying pBOMBL12CRia(*ct226)*-*ct226-3XFLAG*, under uninducing conditions, inclusions demonstrated both FLI1 and LRRF1 localized to the inclusion membrane. Complementation of Ct226-3XFLAG by aTc demonstrates robust localization of both FLI1 and LRRF1 to the inclusion. (B) For the strain carrying pBOMBL12CRia(*ct226*)-*ct225-3XFLAG*, uninducing conditions demonstrated localization of LRRF1, but not FLI1. Complementation of Ct225-3XFLAG did not restore localization of either host protein to the inclusion membrane in induced inclusions. (C) For the strain carrying pBOMBL12CRia(*ct226*)*-ct224-3XFLAG*, FLI1 is not localized to the inclusion membrane in the uninduced condition, while LRRF1 does localize. Neither FLI1 nor LRRF1 localize to the inclusion following complementation of Ct224-3XFLAG.

During our initial characterization of our single complement strains, we observed that induction of the Ct226-3XFLAG complement strain resulted in smaller inclusions. Therefore, we quantified inclusion area to determine if complementation of individual Incs in the pBOMBL12CRia(*ct226*) background altered inclusion size, as quantified by ImageJ. Neither E.V. strain nor L2/*ct226* KD strain demonstrated a difference in inclusion size between uninducing and inducing conditions. Therefore, neither dCas12 expression or simultaneous knockdown of *ct226, ct225,* or *ct224* altered inclusion growth. However, single complementation of Ct226-3XFLAG, in the absence of Ct225 and Ct224, decreased inclusion area by 77% (Figure 3J). Complementation of Ct225-3XFLAG also had a moderate impact on inclusion area (37% decrease), while complementation of Ct224-3XFLAG had no impact on inclusion area (Figure 3J). This suggests that complementation of Ct226 in the absence of Ct225 and Ct224 greatly reduces inclusion size.

### Induction of knockdown in L2/*ct226* KD and L2/*ct225* KD, but not L2/*ct224* KD, negatively impacts FLI1 and LRRF1 localization to the inclusion membrane

To test whether *ct226* knockdown impacts localization of either FLI1 or LRRF1 to the inclusion membrane, HEp2 cells were infected with the indicated knockdown strain or the empty vector control, and knockdown was induced as described as above. Cells were fixed at 24hpi and processed for IFA as described in Materials and Methods. dCas12 expression alone in the EV control strain did not impact localization of FLI1 or LRRF1 to the inclusion membrane (Supplemental Figure 6). For the L2/*ct226* KD strain, both FLI1 and LRRF1 localized to the inclusion membrane under non-inducing conditions (Figure 4A), as seen in wild-type inclusions. However, when knockdown was induced by addition of aTc, neither FLI1 nor LRRF1 localized to the inclusion membrane; instead, both proteins remained in the cytoplasm of the host cells (Figure 4A). This suggests that simultaneous knockdown of *ct226, ct225,* and *ct224* leads to loss of inclusion membrane localization of FLI1 and LRRF1. Induction of the L2/*ct225* KD strain resulted in a statistically significant decrease in fluorescence intensity of both FLI1 (****p-value <0.0001) and LRRF1 (**p-value=0.0011) at the inclusion (Figure 4B & 4D). This suggests that while the localization of both FLI1 and LRRF1 are negatively impacted following knockdown of *ct225*, FLI1 recruitment is more severely affected. Lastly, induction of the L2/*ct224* KD strain did not alter FLI1 or LRRF1 localization to the inclusion (Figure 4C). Overall, these data indicate that knockdown of *ct224* alone does not impact localization, but FLI1 and LRRF1 localization are completely lost in L2/*ct226* KD and greatly reduced in the L2/*ct225* KD strain. These data suggest that simultaneous knockdown of *ct226* and *ct225* results in total loss of both proteins at the inclusion.

### Individual complementation of Ct226-3XFLAG fully rescues localization of both LRRF1 and Flightless 1

We tested if complementation of any of the individual genes restored localization of FLI1 and/or LRRF1 as described above for the knockdown strains. In the strain carrying pBOMBL12CRia(*ct226)-ct226-3XFLAG,* both FLI1 and LRRF1 localized at the inclusion membrane under non-inducing conditions. Under inducing conditions, complementation of Ct226-3XFLAG fully rescued FLI1 and LRRF1 localization (Figure 5A), indicating that Ct226, in the absence of Ct225 and Ct224, is sufficient to restore localization of FLI1 and LRRF1.

We also tested localization of FLI1 and LRRF1 following single complementation of either Ct225-3XFLAG or Ct224-3XFLAG. HEp2 cells were infected with either the Ct225-3XFLAG complementation strain or the Ct224-3XFLAG complementation strain, induced or not as described in the Materials and Methods, and fixed at 24hpi. Coverslips were processed for IFA as described above to visualize FLI1 or LRRF1. Under non-inducing conditions, FLI1 did not localize to the inclusion in the Ct225 and Ct224 complementation strains (Figure 5B & 5C). This was unexpected since we anticipated FLI1 would localize to the inclusion as observed in the L2/*ct226* KD strain under non-inducing conditions (Figure 4A). LRRF1, on the other hand, did localize to the inclusion under non-inducing conditions for both strains (Figure 5B & 5C). Complementation with Ct225-3XFLAG or Ct224-3XFLAG did not restore FLI1 or LRRF1 localization (Figure 5B & 5C), suggesting that neither protein is sufficient to restore FLI1 or LRRF1 recruitment when individually complemented.

### In the absence of LRRF1, complementation of Ct226-3XFLAG rescues FLI1 localization to the inclusion membrane

To definitively test if FLI1 localizes to the inclusion independently of LRRF1, we tested FLI1 localization in the cells infected with the Ct226-3XFLAG complementation strain following LRRF1 siRNA knockdown. HEp2 cells were treated with either non-targeting (NT) or LRRF1-targeting siRNA then infected with the complement strain, subsequently induced, and cells were fixed at 24hpi. LRRF1 knockdown was confirmed by both IFA and western blot (Supplemental Figure 7). Following induction of knockdown and complementation of Ct226-3XFLAG, FLI1 localized to the inclusion membrane in cells treated with either NT or LRRF1 siRNA (Figure 6). This suggests that complementation of Ct226-3XFLAG rescues FLI1 localization to the inclusion membrane in the absence of LRRF1. Since FLI1 requires LRRF1 to co-immunoprecipitate with Ct226 (Figure 4A), this strongly suggests that FLI1 has an alternative mechanism of recruitment to the inclusion membrane that is distinct from LRRF1-mediated recruitment by Ct226.

**Figure 6.**
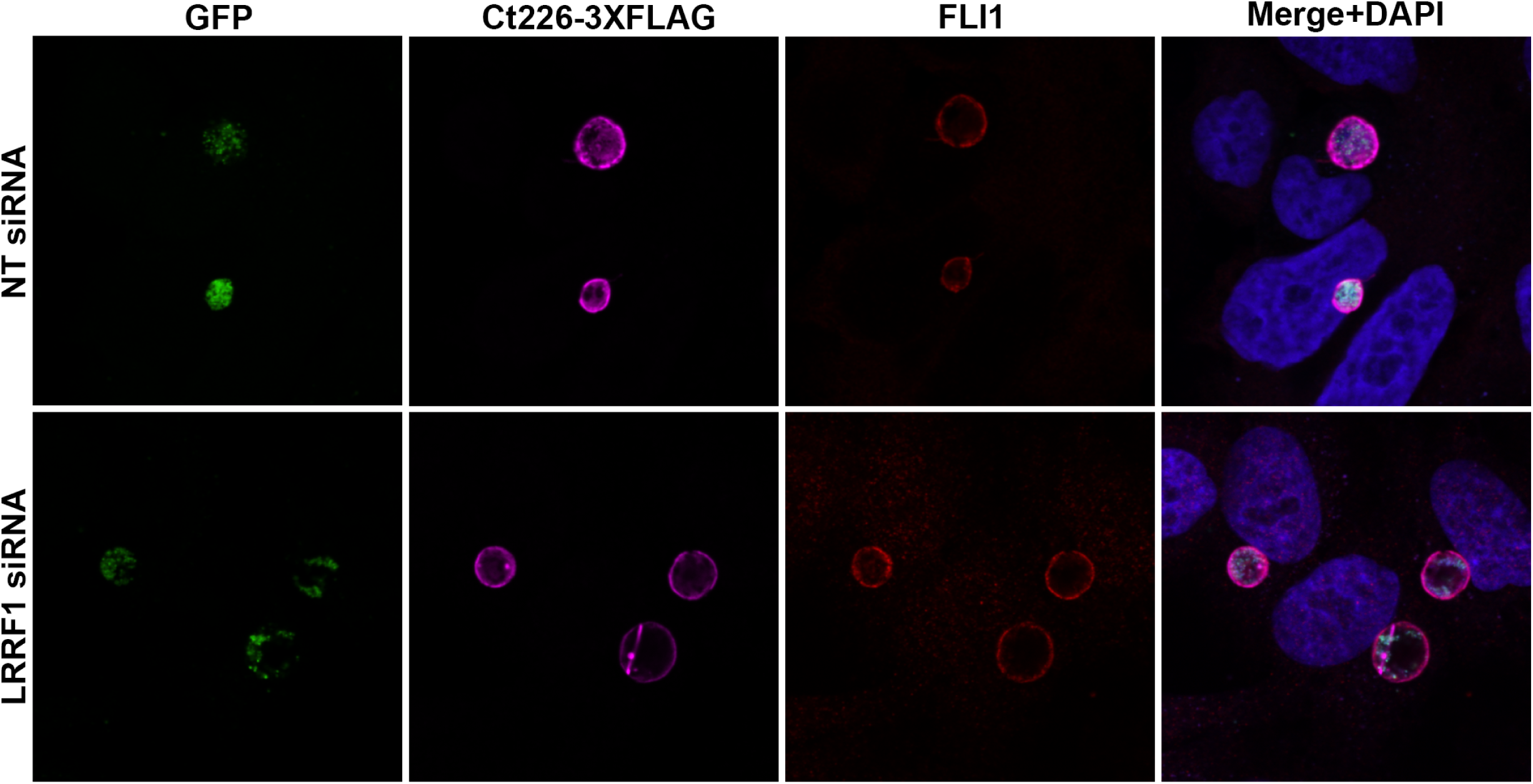
In the absence of LRRF1, complementation of Ct226-3XFLAG rescues FLI1 localization to the inclusion membrane. HEp2 cells were treated with either non-targeting (NT) or LRRF1 siRNA and then infected with the Ct226-3XFLAG complementation strain. Induction of dCas12 and Ct226-3XFLAG expression was induced as described in Materials and Methods and cells were fixed at 24hpi by 4% paraformaldehyde. Immunofluorescence was performed to visualize chlamydial organisms (GFP; green), Ct226-3XFLAG (anti-FLAG; magenta), and FLI1 (anti-FLI1; red). Host and bacterial DNA were visualized with DAPI (blue). Ct226-3XFLAG complementation restored FLI1 localization to the inclusion in both NT and LRRF1 siRNA-treated cells.

## DISCUSSION

Of all chlamydial effector proteins, candidate *inc* genes make up an estimated 7% of open reading frames within the highly reduced chlamydial genome (63), highlighting their importance to chlamydial pathogenesis. Building on our previous studies (28), we sought to describe the Inc-host protein interactions required for recruitment of FLI1 and LRRF1—two eukaryotic proteins that are involved in multiple cell-signaling networks and regulatory processes. In this work, we report the dynamics of FLI1 and LRRF1 localization to the inclusion membrane and identify the chlamydial proteins involved in their recruitment. We demonstrated FLI1 localizes to the inclusion membrane between 12 and 14hpi, (early mid-cycle of chlamydial development), and it remains at the inclusion through 46hpi, which aligns closely with the timeline described for LRRF1 localization (28). We hypothesize that the stable localization at the inclusion means these proteins are either i) being sequestered away from their native functions, or ii) their functions are being relocated to the inclusion membrane. Understanding how these proteins localize to the inclusion is an important first step towards how to investigate their function.

We demonstrate that both the LRR and gelsolin domains of FLI1 are capable of localizing to the inclusion, however, FLI1 localization driven by the LRR domain is more efficient. We also demonstrated FLI1 interacts with Ct226 in complex with LRRF1, and while FLI1 depends on LRRF1 for interaction with Ct226, both proteins can localize to the inclusion independently of each other. Together, these data suggest multiple mechanisms for FLI1 localization aside from interaction with Ct226 and LRRF1.

Of note, two other ongoing studies in the field have implicated Ct226 in FLI1 recruitment to the inclusion in a LRRF1-dependent manner. Interestingly, these studies from the Lutter group and Engel group use different genetic methods to knock-out (KO) *ct226* in contrast to our inducible CRISPRi knockdown approach. For example, the Lutter group has generated a *ct226* knockout strain using type II intron mutagenesis (i.e. TargeTron) (C. Holcomb, E. Lutter; personal communication), while the Engel team has created a *ct226* knockout strain using fluorescence-reported allelic exchange mutagenesis (FRAEM) (C. Elwell, J. Engel; personal communication). Indeed, some of our results agree with their findings. For example, we demonstrated that simultaneous knockdown of *ct226*, *ct225*, and *ct224* resulted in a complete loss of localization of either FLI1 or LRRF1 during chlamydial infection and only individual complementation of Ct226 in this background fully restored FLI1 and LRRF1 localization. These data would suggest that expression of Ct226 alone leads to FLI1 localization to the chlamydial inclusion. However, we were unable to demonstrate an interaction between Ct226 and FLI1 via co-immunoprecipitation unless LRRF1 was present. In addition, we observed that FLI1 can localize to Ct226-positive (i.e., wild-type) inclusions in the absence of LRRF1 (Figure 2A). Similarly, expression of Ct226-3XFLAG following knockdown of *ct226* in LRRF1 siRNA-treated cells resulted in recruitment of FLI1 to the inclusion (Figure 6). Taken together, these data are strong evidence for an alternative, LRRF1-independent recruitment mechanism for FLI1 that does not rely on a direct protein-protein interaction with Ct226. Instead, it may be that Ct226 interacts with another Inc that recruits FLI1 to the inclusion membrane. Importantly, our FLI1 truncation studies (Figure 1B-D) indicate this interaction could be mediated either via its LRR or gelsolin domains. Overall, it will be important to assess if these other genetic models lead to polar effects in downstream genes, particularly for the TargeTron approach, to clarify the function of Ct226. Our findings also highlight the advantage of the CRISPRi inducible knockdown system to investigate the impact of knockdown of multiple gene targets that led us to look further into the interactions between these chlamydial proteins.

It is intriguing that simultaneous knockdown of three genes (*ct226, ct225,* and *ct224*) in the L2/*ct226* KD strain did not alter inclusion area, but individual complementation of Ct226-3XFLAG expression with 2nM aTc in this genetic background produced inclusions that were 77% smaller in area compared to uninducing conditions (Figure 3J). We previously showed that overexpression of Ct226-FLAG with 5nM aTc in Ctr L2 led to a 25% decrease in inclusion area (28). These data indicate that restriction of growth due to Ct226 overexpression is limited when there are endogenous levels of Ct225 and Ct224 expression. It is possible that Ct226 cooperates with Ct225, Ct224, or other Incs for optimal inclusion growth. Thus, overexpression of Ct226 in the absence of the other proteins may concentrate other components of the inclusion membrane and/or disrupt canonical Inc-host and Inc-Inc interactions. To test these hypotheses, combining genetic knockdown with multiplexed complementation of two or more genes would help determine which Incs functionally cooperate with each other during chlamydial development. Additionally, co-infection models using strains expressing Incs tagged with different epitopes could also be used to determine interactions between Ct226, Ct225, and Ct224 (64, 65).

The localization of Ct225 to the bacterial membrane, as opposed to its anticipated localization to the inclusion membrane, was an unexpected finding in this study. Ct225 is the smallest member of the *ct227* operon (∼13-15 kDa) and was annotated as an Inc in the very first bioinformatics screening using the conserved bilobed transmembrane domain to identify putative Incs in *C. trachomatis* (20). It was identified again in another study that expanded this same bioinformatics screen to seven species of *Chlamydiae* (66). Ct225 localization was first mentioned in a study measuring temporal expression of chlamydial genes, saying Ct225 had been “detected in association with the inclusion membrane by immunofluorescent staining,” but the data were not shown (16). In 2008, Li and colleagues (61) characterized 50 putative inclusion membrane proteins by raising antibodies against chlamydial proteins fused to GST. Indirect immunofluorescence studies using anti-GST-Ct225 antibodies localized Ct225 to the chlamydial inclusion membrane in *C. trachomatis* infected HeLa cells (this same antibody was generously shared and used for data shown in Supplemental Figure 5) (67). Once chlamydial transformation protocols were developed, a study published by Weber et al. in 2015 expressed predicted Incs fused to FLAG tags to determine their localization by immunofluorescence microscopy. However, while Ct225 was included in the study design, it was annotated as ‘not expressed’ (62). Therefore, our study is the first to confirm ectopic expression of Ct225 from a chlamydial plasmid and localization of endogenous and ecopicaly expressed Ct225 by immunofluorescence microscopy.

Because our data demonstrate that Ct225 does not localize to the inclusion membrane, it is interesting that our studies showed induction of knockdown in the L2/*ct225* KD strain led to a statistically significant decrease in FLI1 and LRRF1 recruitment to the inclusion membrane, despite unaffected Ct226 protein levels (Figure 4B & 4D; Supplemental Figure 3). Since Ct225 is situated in the bacterial membrane and can interact with Ct226 by bacterial adenylate cyclase two-hybrid assay (BACTH) (Supplemental Figure 8), Ct225 may be important for efficient type III secretion of Ct226. Hence, knockdown of *ct225* might lead to decreased Ct226 in the inclusion membrane and subsequently decreased recruitment of host proteins LRRF1 and FLI1. However, the Ct226-3XFLAG complement strain offers evidence against the interpretation that Ct225 is *required* for Ct226 secretion since Ct226-3XFLAG is secreted and found within the inclusion membrane even while *ct225* is knocked down (Figure 3G). As such, it is possible that Ct225 serves a positive regulatory function for Ct226 secretion, or, alternatively, it may be involved in organization and stability of Ct226 in the inclusion membrane. Additional studies are needed to clarify the function and molecular interactions of Ct225 during *C. trachomatis* infection.

The goal of this study was to better understand the mechanisms of FLI1 and LRRF1 recruitment to the inclusion by Inc proteins. We established that FLI1 recruitment occurs during early mid-cycle development and interacts in complex alongside Ct226 and LRRF1. We also present evidence that FLI1 has a recruitment mechanism independent of LRRF1. Both observations are critical for identifying which host signaling pathways are impacted by FLI1/LRRF1 localization and for determining how *C. trachomatis* might be modifying them to create an optimal intracellular niche. To this end, the multitude of strains developed in this study will be used to test the impact of FLI1/LRRF1 localization on host cell processes. Importantly, we have shown the adaptability of the CRISPRi knockdown system to investigate Inc-host, as well as Inc-Inc, interactions.

Understanding these complex interactions and how they change throughout development will help us to understand how *C. trachomatis* manipulates its environment to establish a successful infection.

## MATERIALS AND METHODS

### Tissue Culture and Chlamydial Strains

HEp2 cells (Ouellette lab stock), HeLa cells (CCL-2.1; American Type Culture Collection [ATCC], Manassas, VA), and McCoy cells (CRL-1696; ATCC, Manassas, VA) were routinely passaged and were cultured in Dulbecco’s Modified Eagle Media (DMEM; Gibco/Thermo Fisher) supplemented with 10% fetal bovine serum (Sigma-Aldrich, St. Louis, MO) and 10 μg/ml gentamicin (Gibco/Thermo Fisher). All cells were incubated at 37°C at 5% CO_2_. *Chlamydia trachomatis* serovar L2 (lymphogranuloma venereum [LGV] strain L2/434/Bu) was propagated using HeLa cells and purified for use in experiments using density gradient centrifugation as described in previous protocols (68). HeLa or McCoy cells were used for LGV 434 transformation of the strains produced in this study. Chlamydial strain titers were determined by measuring by number of inclusion forming units (IFUs), using previously described methods (69). HeLa cells were used to determine titers and titers were used to determine multiplicity of infection (MOI) for subsequent experiments. For all strains, cells were infected at the indicated MOI by centrifugation at 400xg for 15 minutes and media was replaced following 15-minute incubation at 37°C at 5% CO_2_. All cell lines and media were routinely tested for *Mycoplasma* spp. contamination (Lookout Mycoplasma PCR detection kit; Sigma-Aldrich, St. Louis, MO).

### Plasmid construction

Sequences for primers and crRNAs are provided in Supplemental Table 1. For the knockdown strains, the vector pBOMBL12CRia::L2 (pBOMBL12CRia) (53) was modified by BamHI-digest, treated with alkaline phosphatase (AP), and the appropriate crRNA gBlock (IDTDNA; Coralville, IA) was inserted using the NEBuilder HiFi DNA assembly kit (New England Biolabs (NEB); Cambridge, MA) according to manufacturer protocols. The resulting plasmids were transformed into chemically competent *E. coli* 10-β using conventional techniques. Plasmids from transformants were isolated by miniprep (Qiagen), screened for the correct plasmid by colony PCR or plasmid digest, and were confirmed by sequencing across the crRNA insert site (Genewiz/Azenta).

For the complementation plasmids, the pBOMBL12CRia(*ct226*) plasmid was modified by SalI-digest and insertion of a gBlock (New England Biolabs (NEB); Cambridge, MA) containing a ribosomal binding site (RBS), a KpnI digest site, and the epitope tag 3XFLAG. Open reading frames (ORFs) for *ct226*, *ct225*, and *ct224* were amplified from *C. trachomatis* L2 genomic DNA using Phusion DNA polymerase (New England BioLabs, Ipswich, MA) and purified using the QIAquick PCR Purification Kit (Qiagen; Hilden, Germany). Purified PCR products were inserted into KpnI-digested pBOMBL12CRia(*ct226)*-3XFLAG vector as described above. The resulting plasmids were isolated as described above and confirmed by sequencing across the insertion site.

### Anhydrotetracycline (aTc) induction conditions for chlamydial strains

For pBOMB4-*ct226-FLAG* and pBOMBLmT-*ct225*-*FLAG*, expression was induced at 7hpi with 5nM and 2nM aTc, respectively. For all knockdown and complement strains, knockdown and/or complementation was induced using 2nM aTc. Induction timeline for the strains is as follows: For L2/*ct226* KD strain and the empty vector control strain, infected cells were induced at 3hpi. For strains L2/*ct225* KD and L2/*ct224* KD, samples were induced at time of infection. For all complement strains, inclusions were induced with 2nM aTc at 7hpi.

### Antibodies and indirect immunofluorescence

Primary antibodies used in these studies were polyclonal rabbit anti-LRRF1 (Bethyl Laboratories), sheep anti-IncA, rabbit anti-FLI1 (Thermo Fisher), goat anti-MOMP (Meridian), mouse anti-FLAG (Sigma-Aldrich), rabbit anti-FLAG (Sigma-Aldrich), mouse anti-Ct225-GST (gift from Guangming Zhong, University of Texas Health Sciences Center-San Antonio), rabbit anti-Ct226 (gift from Erika Lutter, Oklahoma State University), mouse anti-AsCpf1 (dCas12; Sigma-Aldrich). Secondary antibodies for indirect immunofluorescence included donkey anti-647, -594, and -488 (Jackson Labs, Bar Harbor, Maine). For indirect immunofluorescence microscopy, antibodies were diluted as indicted in 3% bovine serum albumin (BSA) in PBS and incubated for 1 hour at 37°C with the exception of Rb anti-FLI1. Rb anti-FLI1 (Invitrogen-Thermo Fisher) was incubated for 2 hours at 4 °C and rocked overnight at 4 °C in DPBS. DNA detection was performed using DAPI (4′,6-diamidino-2-phenylindole). Western blots were visualized by secondary antibodies conjugated to IRDye 680 or IRDye 800 (LiCor Biosciences, Lincoln, NE).

### Chlamydial transformation

Chlamydial transformations in this study were performed as described previously (67). Briefly, either HeLa or McCoy cells were used for chlamydial transformation and were seeded in 6-well plates at a density of 10^6^ on the day prior. 2ug of plasmid was added to purified EBs in a Tris-CaCl_2_ buffer solution and allowed to incubate for 30 min at room temperature. Following incubation, the transformation mix was diluted in Hanks’ balanced salt solution (HBSS; Gibco) and added to one well of the 6-well plate. Plates were centrifuged at 400xg for 15 minutes and media was replaced with DMEM+ 10% FBS following 15 min of incubation at 37°C at 5% CO_2_. At 8hpi, penicillin was added at either 1U/ml or 2U/ml, and cycloheximide was added at concentration of 1ug/mL (Millipore Sigma). At 48hpi, infected cells were harvested and centrifuged at 17,000 x g for 30 min at 4°C. Pellets were resuspended in 1mL of HBSS and centrifuged for 5 min at 400 x g at 4°C. The supernatant containing the infectious progeny was added to a fresh monolayer of either HeLa or McCoy cells seeded the day before. This cycle of infection followed by progeny harvest was continued until only wild-type inclusions constitutively expressing GFP from the transformed plasmid were observed. Transformed EBs were collected in 2-sucrose-phosphate (2SP) for storage at -80°C and future plaque-cloning and expansion.

### Inclusion forming unit (IFU) assay

Inclusion forming unit assays, or infectious progeny assays, were performed as previously described (70, 71). Briefly, all knockdown and complement strains were used to infect a monolayer of HeLa cells in a 24-well plate. Strains were either induced using anhydrotetracycline (aTc) or not, as described above. At 24hpi, infected cells were scraped and vortexed with 1-mm glass beads, and frozen at -80°C. Lysates were collected and used to infect a new monolayer of HeLa cells. Inclusions were counted and reported as IFUs per ml. A paired student’s t analysis was performed to determine significant differences between uninduced and induced conditions for each strain.

### Time-course localization of Flightless 1 to the inclusion membrane

HEp2 cells were plated in 24-well plates containing cell culture-treated coverslips and infected with WT *Chlamydia trachomatis* serovar L2 (MOI=1). Coverslips were fixed with 4% paraformaldehyde at the indicated time points post infection and permeabilized with 0.5% TritonX-100. Coverslips were processed for indirect immunofluorescence to visualize endogenous FLI1 (green), chlamydial organisms (MOMP; red), and DNA (DAPI; blue).

### Eukaryotic cell transfection

HEp2 cells were plated on coverslips placed in wells of 24-well plates. 24 hours after plating, 850ng of the indicated construct was transfected using JetPrime transfection reagent (Polyplus; Illkirch-Graffenstaden, France) according to manufacturer protocol. Cells were returned to the incubator at 37°C at 5% CO_2_. After 4 hours of incubation with the transfection media, media was replaced with fresh DMEM+10% FBS and allowed to rest for 2 hours. Cells were then infected (MOI=1) with wild-type Ctr L2, fixed at 24hpi with 4% paraformaldehyde, and process for immunofluorescence as described above.

### siRNA knockdown of LRRF1 and Flightless 1 and co-immunoprecipitation of Ct226-FLAG following

Nontargeting siRNA (catalog number SR30004; Origene, Rockville, MD), pooled LRRF1 siRNA (catalog numbers 43450, s229968, and s17599; Ambion Life Technologies), and FLI1 (catalog number L-017506-01-0010; Dharmacon, Lafayette, CO) were used in knockdown experiments. HEp2 cells were seeded in six-well plates at a density of 1x10^6^ cells/well and 24 hours post-plating, cells were transfected with 60nM the appropriate siRNA using JetPrime Transfection Reagent (Polyplus), according to manufacturer protocols. 24 hours after siRNA treatment, media on the cells was replaced and 48 hours following siRNA transfection, cells were infected with pBOMB4-Ct226-FLAG strain (MOI=1) and induced as described above (28). Cell lysates were harvested 24hpi using RIPA buffer (50mM Tris-HCl; pH 7.4, 150nM NaCl, 0.1% SDS, and 0.5% sodium deoxycholate) amended with 1% TritonX-100, 1X HALT Protease Inhibitor Cocktail (Thermo Fisher Scientific, Waltham, MA), 1X universal nuclease (Pierce, Rockford, IL), and 150uM CPAF inhibitor, clasto-lactacystin β-lactone (Santa Cruz Biotechnology, Dallas, TX). Solubilized lysates were clarified, and protein concentration was determined by protein assay (EZQ Protein Quantification Kit; Life Technologies, Carlsbad, CA). Ct226-FLAG was affinity purified using magnetic FLAG beads (Sigma-Aldrich, St. Louis, MO). Bound proteins were eluted by addition of 500ug/mL FLAG peptide (Thermo Fisher) in 50uL of RIPA+1% TritonX-100.The eluate fraction was combined with equal volume of 2X Laemelli amended with 5% β-mercaptoethanol.

Samples were resolved by running on a 8% polyacrylamide gel, transferred to polyvinylidene difluoride (PVDF) membrane (pore size, 0.45 μm; Thermo Fisher) and blotted for FLI1 (144 kDa), LRRF1 (dimer; 160 kDa), and Ct226-FLAG (∼19 kDa).

### Indirect Immunofluorescence microscopy for strain characterization and localization studies

HEp2 cells were plated on coverslips placed in a 24-well plate with three replicate wells to visualize dCas12, FLI1, or LRRF1. Cells infected with the appropriate strain (MOI=0.8) and induced as described above. At 24hpi, cells were fixed using 4% paraformaldehyde and permeabilized with 0.5% Triton-X 100. Chlamydial organisms (green) were visualized by constitutive GFP expression from the pBOMBL plasmid. Host and chlamydial DNA was visualized by DAPI (blue). dCas12, LRRF1, or FLI1 were visualized in the red. Coverslips were mounted on slides using ProLong Glass Antifade mounting medium (Thermo Fisher) and imaged using a Zeiss ApoTome.2 fluorescence microscope at 100x magnification.

### RT-qPCR analysis of knockdown and complementation strains

HEp2 cells were plated in 6-well plates at a density of 1x10^6^, and 24 hours post-plating were infected (MOI=1) with the appropriate strain and induced as described above. RNA was collected at 3, 12, and 24hpi using Trizol (Invitrogen/Thermo Fisher) and chloroform extraction was performed for RNA isolation. DNA contamination was removed by use of Turbo DNAfree (Ambion/Thermo Fisher) and done in accordance with manufacturer protocols. DNAse-treated RNA was converted to cDNA by incubation of SuperScript III Reverse Transcriptase (Invitrogen/Thermo Fisher) with random nonamers (New England Biolabs, Ipswich, MA). Resulting cDNA was diluted and stored at -80°C. Diluted cDNA was used in equal volume with SYBR green master mix (Applied Biosystems) for qPCR. Transcripts for *ct226, ct225*, *ct224, ct227*, *ct223* and *16s* were quantified using the standard amplification cycle with a melting curve analysis measured by QuantStudio 3 (Applied Biosystems/Thermo Fisher). A complete list of qPCR primer pairs is provided in Supplemental Table 1. A standard curve of genomic DNA was generated from isolated *C. trachomatis* serovar L2 genomic DNA. Transcripts were normalized to *16s* transcript levels.

### Quantification of inclusion area

HEp2 cells were infected with the indicated strains (MOI=1). Cells were fixed at 24hpi and stained for immunofluorescence as described above. ImageJ was used to quantify inclusion area using the freehand selection tool. The area of 40 inclusions was quantified per condition for each strain. An ordinary two-way ANOVA test was used to determine statistical differences.

### Quantification of fluorescence intensity at the chlamydial inclusion membrane for L2/*ct225* KD strain

HEp2 cells were infected with the L2/*ct225* KD and processed for indirect immunofluorescence microscopy as described above for the FLI1 and LRRF1 localization studies. Exposure time was maintained for both the uninduced and induced conditions. Quantification of fluorescence was performed using ImageJ. Briefly, the “Free Hand Line” tool was used to manually draw lines around the inclusion membranes on the “merged channel” image. “Line Width” was set to 12 to cover the width of the inclusion membrane. Once lines were drawn for inclusions on the “merged channel” image, the image for the channel visualizing FLI1 or LRRF1 was used to measure raw integrated density. Data were collected for 90 inclusions for uninduced and induced samples. Because smaller inclusions have a smaller surface area, fluorescence intensity of proteins localized at the inclusion is greater than for larger inclusions. To account for this, data were normalized to inclusion perimeter and reported as raw integrated density/inclusion perimeter. An unpaired student’s t-test was performed to determine statistical significance between uninduced and induced inclusions.

### Bacterial adenylate cyclase two-hybrid (BACTH) and β-Galactosidase assays

The BACTH assay was used to test interactions between Ct226, Ct225, and Ct224. The basis for this assay is two genes of interest are translationally fused to either the T25 or T18 subunits of *Bordetella pertussis* adenylate cyclase. If the proteins interact, it results in functional reconstitution of cyclic AMP (cAMP) production in DHT1 *E. coli* lacking adenylate cyclase (Δ*cya*) and drives expression of β-galactosidase under the *lac* promoter. This assay was performed as previously described (21, 72, 73). Briefly, pST25 and pUT18C vectors fused to either IncA, Ct226, Ct225, or Ct224 (Supplemental Table 1) were co-transformed into chemically competent DHT *E. coli* (Δ*cya*) and were plated on minimal medium M63 selection plates containing IPTG, 40μg/ml 5-bromo-4-chloro-3-indolyl-β-D-galactopyranoside (X-Gal), 0.04%casein hydrolysate, and 0.2% maltose. Homotypic interactions of IncA were used for the positive control while PST25 fused to Ct226 co-transformed with the pUT18C empty vector was used as the negative control. Blue colonies were indicative of a positive interaction and representative plates were imaged (Supplemental Figure 8A). For the β-galatosidase assay, random colonies were picked and grown in M63 minimal media prior to permeabilization with chloroform and 0.1% SDS, which extracts the β-galactosidase. The reaction was allowed to proceed for exactly 20 min, at which it was halted by addition of 1 M NaHCO_3_ and 0.1% *o*-nitrophenol-β-galactoside (ONPG). Absorbance at the 405 nm wavelength was measured by a Tecan plate reader, normalized to optical density (bacterial growth), and β-galactosidase activity reported as relative units (RU).

## Data availability

All original data that contributed to the final figures will be uploaded into Mendeley. Upon acceptance this section will be updated with a digital object identifier to indicate where the original data can be found.

## ACKNOWLEDGEMENTS

We would like to thank Guangming Zhong for his gift of the endogenous Ct225 antibody. We would also like to thank Erika Lutter for her gift of the Ct226 endogenous antibody. Lastly, we are deeply appreciative of the conversations and critical feedback we received from Joanne Engel, Cherilyn Elwell, Erika Lutter, Christian Holcomb and other members of their respective laboratories. We sincerely thank Lindsey Knight for her technical assistance during these studies. These studies were supported by UNMC Start-up funds to EAR and a UNMC Graduate Fellowship to NAS.

## FIGURE LEGENDS

**Supplemental Figure 1. Control for siRNA knockdown of FLI1 or LRRF1 by indirect immunofluorescence.** Indirect immunofluorescence was used to confirm siRNA knockdown of FLI1 or LRRF1. (A) FLI1 visualization at the inclusion membrane following treatment with either non-targeting (NT) siRNA or FLI1 siRNA. FLI1 localizes to the inclusion membrane in cells treated with NT siRNA. It is not visible at the inclusion following treatment with FLI1 siRNA. (B) LRRF1 visualization at the inclusion membrane following treatment with either NT siRNA or LRRF1 siRNA. LRRF1 localizes to the inclusion membrane in cells treated with NT siRNA. It is not visible by IFA following treatment with LRRF1 siRNA.

**Supplemental Figure 2. Representative western blot demonstrating co-immunoprecipitation of Flightless 1 with *C. trachomatis* L2 expressing Ct226-FLAG.** HEp2 cells were seeded, infected with the strain carrying pBOMB4-*ct226-FLAG* and induced or not, according to the Materials and Methods. Cell lysates were collected at 24hpi, Ct226-FLAG was affinity purified, eluate fractions were separated by 8% SDS-PAGE gel and transferred to 0.45nm PVDF membrane. The resulting western blot was blotted for FLI1 or LRRF1 and Ct226-FLAG (anti-FLAG) was used as the loading control. Ct226-FLAG was only detected in the induced samples. (∼19.2 kDa). LRRF1 (dimer; 160 kDa), used a positive control for Ct226-FLAG pulldown, was detected in the eluate fraction of induced samples. FLI1 (144 kDa) was also detected in the eluate fraction of induced samples. Data represent two technical replicates and are representative of three biological replicates.

**Supplemental Figure 3. Detection of endogenous *Ct226* protein in L2/*ct226* KD and L2/*ct225* KD strains.** To determine if endogenous Ct226 protein is decreased in the knockdown strains targeting either *ct226* or *ct225*, HEp2 cells were infected with the indicated strain and induced or not. Whole cell lysates were collected at 24hpi, clarified, and blotted for endogenous Ct226 (anti-Ct226; gift from Erika Lutter) and chlamydial MOMP (anti-MOMP) was used as a loading control. For the L2/*ct225* KD strain, Ct226 (∼19 kDa) was detected in both uninduced and induced samples. For the L2/*ct226* KD strain, endogenous Ct226 protein levels were decreased in induced samples compared to uninduced samples.

**Supplemental Figure 4. Detection of 3XFLAG-tagged protein in complement strains by western blot.** To assess the expression level of the complemented proteins in each of our complementation strains in both uninduced and induced conditions, protein samples were collected and assessed by western blot. HEp2 cells were plated, infected with one of the complement strains, and induced as described in the Materials and Methods. Whole cell lysates were blotted with anti-FLAG antibody to detect the complemented protein, as indicated by the asterisk (Ct226-3XFLAG, ∼19.2 kDa; Ct225-3XFLAG, ∼13-15 kDa; Ct224-3XFLAG, ∼16 kDa). For complementation of Ct226-3XFLAG, there was evidence of leaky expression of Ct226 in the uninduced sample and was expressed robustly following addition of aTc. Ct225-3XFLAG was detected only in the induced sample at the predicted weight. Ct224-3XFLAG was not detectable in uninduced sample but was detectable following addition of aTc at the appropriate weight.

**Supplemental Figure 5. Ct225 localization in wild-type *C. trachomatis* and strain expressing Ct225-FLAG using endogenous Ct225 antibody.** (A-B) Ectopic expression of Ct225-FLAG at 17 and 24hpi. HEp2 cells were infected with strain carrying pBOMBLmT-Ct225-FLAG and induced with either 0, 0.1, or 0.5 nM aTc at 7hpi. Cells were fixed with methanol at 17hpi and 24hpi and indirect immunofluorescence was used to visualize Ct225-FLAG (anti-FLAG; red), chlamydial organisms (anti-MOMP; green), and host and chlamydial DNA (DAPI; blue). (A) At 17hpi, Ct225-FLAG was visible at both 0.1 and 0.5 nM aTc level of induction and localized to the bacterial membrane. (B) Ct225-FLAG remained localized to the bacterial membrane at 24hpi. (C) Localization of endogenous Ct225. To determine the localization endogenous Ct225, we used a Ct225 endogenous antibody raised against a Ct225-GST fusion (kind gift of Dr. Guangming Zhong) and used it to localize Ct225 in *C. trachomatis* serovar L2 inclusions (anti-Ct225; red channel, alongside chlamydial organisms (anti-MOMP; green) and DNA (DAPI; blue). (D) We used the same anti-Ct225 antibody to determine if the antibody recognized the exogenously expressed Ct225-FLAG construct. Indirect immunofluorescence was used to visualize endogenous Ct225 (anti-Ct225; green), Ct225-FLAG (anti-FLAG; red), and DNA (DAPI). Images were taken using a Zeiss ApoTome.2 fluorescence microscope at 100x magnification. Scale bar= 2μm.

**Supplemental Figure 6. LRRF1 and FLI1 localization in the empty vector control strain.** To ensure that expression of dCas12 did not affect LRRF1 and FLI1 localization during chlamydial infection, we used indirect immunofluorescence to localize FLI1 and LRRF1 during infection with the empty vector control strain carrying the pBOMBL12CRia(*E.V.*) plasmid under both uninducing and inducing conditions. (A) FLI1 localized to the inclusion in both in uninduced and induced conditions. (B) LRRF1 also localized to the inclusion regardless of induction condition. Images were taken using a Zeiss ApoTome.2 fluorescence microscope at 100x magnification. Scale bar= 2μm.

**Supplemental Figure 7. Confirmation of LRRF1 siRNA knockdown by immunofluorescence and western blot in cells infected with the Ct226-3XFLAG complement strain.** HEp2 cells were plated on coverslips and treated with either non-targeting (NT) or LRRF1 siRNA and then infected with the strain carrying pBOMBL12CRia(*ct226*)-*ct226-3XFLAG*. Induction of dCas12 and Ct226-3XFLAG expression was induced or not as described in Materials and Methods and cells were fixed at 24hpi by 4% paraformaldehyde. Replicate wells were collected for protein samples to confirm LRRF1 knockdown by western blot. (A) Indirect immunofluorescence was used to visualize chlamydial organisms (GFP; green), the inclusion membrane (anti-IncA; magenta), and LRRF1 (anti-FLI1; red) as described in Materials and Methods. Host and bacterial DNA were visualized with DAPI (blue). Images are representative inclusions from the induced condition. In cells treated with NT siRNA, LRRF1 localizes to the inclusion. LRRF1 is undetected in cells treated with LRRF1 siRNA. (B) A representative western blot detection of LRRF1 in NT and LRRF1 siRNA treated cells. Whole cell lysates were collected at 24hpi, resolved by SDS-PAGE, and transferred to a PVDF membrane for western blotting. The membrane was probed for LRRF1 (anti-LRRF1; 160 kDa dimer) and GAPDH was used as a loading control (anti-GAPDH; 36 kDa). In NT siRNA treated cells, the LRRF1 dimer is detected at the appropriate molecular weight (correct indicated by the left facing arrow) while LRRF1 is not detectable in LRRF1 siRNA treated samples. The experiment included two technical replicates as indicated by n1 and n2 labeled brackets.

**Supplemental Figure 8. Interaction of Ct226 with other candidate Incs in the *ct227* gene cluster by bacterial adenylate cyclase two-hybrid (BATCH) assay.** To determine if Ct226 can interact with Ct225 or Ct224, we used a bacterial adenylate cyclase two-hybrid (BACTH) assay followed by a beta-galactosidase assay to both qualitatively and quantitatively assess protein-protein interactions. Ct226 fused to the T25 fragment of *Bordetella pertussis* adenylate cyclase was co-transformed with Inc fusions to the T18 fragment adenylate cyclase. IncA homotypic interactions were used as a positive control and co-transfection with the T18-empty vector was used as a negative control. (A) Representative colonies indicating interactions. (B) Quantitative analysis of interactions measured by beta-galactosidase assay and reported as relative units. Values five times the negative control are considered positive. Ct226 demonstrated strong homotypic interactions, which has been previously reported (28). It also demonstrated interaction with Ct225, but not with Ct224.

## References

1. Rodrigues R, Sousa C, Vale N. 2022. Chlamydia trachomatis as a Current Health Problem: Challenges and Opportunities. Diagnostics (Basel) 12.

2. Anonymous. Centers for Disease Control and Prevention. Sexually Transmitted Disease Surveillance 2020. Atlanta: U.S. Department of Health and Human Services; 2022. .

3. Elwell C, Mirrashidi K, Engel J. 2016. Chlamydia cell biology and pathogenesis. Nature Reviews Microbiology 14:385–400.

4. Geisler WM, Suchland RJ, Whittington WL, Stamm WE. 2003. The relationship of serovar to clinical manifestations of urogenital Chlamydia trachomatis infection. Sex Transm Dis 30:160–5.

5. Detels R, Green AM, Klausner JD, Katzenstein D, Gaydos C, Handsfield H, Pequegnat W, Mayer K, Hartwell TD, Quinn TC. 2011. The incidence and correlates of symptomatic and asymptomatic Chlamydia trachomatis and Neisseria gonorrhoeae infections in selected populations in five countries. Sex Transm Dis 38:503–9.

6. Carey AJ, Beagley KW. 2010. Chlamydia trachomatis, a hidden epidemic: effects on female reproduction and options for treatment. Am J Reprod Immunol 63:576–86.

7. den Heijer CDJ, Hoebe C, Driessen JHM, Wolffs P, van den Broek IVF, Hoenderboom BM, Williams R, de Vries F, Dukers-Muijrers N. 2019. Chlamydia trachomatis and the Risk of Pelvic Inflammatory Disease, Ectopic Pregnancy, and Female Infertility: A Retrospective Cohort Study Among Primary Care Patients. Clin Infect Dis 69:1517–1525.

8. Menon S, Timms P, Allan JA, Alexander K, Rombauts L, Horner P, Keltz M, Hocking J, Huston WM. 2015. Human and Pathogen Factors Associated with Chlamydia trachomatis-Related Infertility in Women. Clin Microbiol Rev 28:969–85.

9. Hillis SD, Owens LM, Marchbanks PA, Amsterdam LF, Mac Kenzie WR. 1997. Recurrent chlamydial infections increase the risks of hospitalization for ectopic pregnancy and pelvic inflammatory disease. Am J Obstet Gynecol 176:103–7.

10. Beatty WL, Byrne GI, Morrison RP. 1994. Repeated and persistent infection with Chlamydia and the development of chronic inflammation and disease. Trends in Microbiology 2:94–98.

11. Bastidas RJ, Elwell CA, Engel JN, Valdivia RH. 2013. Chlamydial intracellular survival strategies. Cold Spring Harb Perspect Med 3:a010256.

12. Abdelrahman YM, Belland RJ. 2005. The chlamydial developmental cycle. FEMS Microbiology Reviews 29:949–959.

13. Hackstadt T, Fischer ER, Scidmore MA, Rockey DD, Heinzen RA. 1997. Origins and functions of the chlamydial inclusion. Trends Microbiol 5:288–93.

14. Hybiske K, Stephens RS. 2007. Mechanisms of host cell exit by the intracellular bacterium Chlamydia. Proc Natl Acad Sci U S A 104:11430–5.

15. Belland RJ, Zhong G, Crane DD, Hogan D, Sturdevant D, Sharma J, Beatty WL, Caldwell HD. 2003. Genomic transcriptional profiling of the developmental cycle of Chlamydia trachomatis. Proc Natl Acad Sci U S A 100:8478–83.

16. Shaw EI, Dooley CA, Fischer ER, Scidmore MA, Fields KA, Hackstadt T. 2000. Three temporal classes of gene expression during the Chlamydia trachomatis developmental cycle. Mol Microbiol 37:913–25.

17. Rucks EA. 2023. Type III Secretion in Chlamydia. Microbiol Mol Biol Rev doi:10.1128/mmbr.00034-23:e0003423.

18. Fields KA, Hackstadt T. 2000. Evidence for the secretion of Chlamydia trachomatis CopN by a type III secretion mechanism. Mol Microbiol 38:1048–60.

19. Bannantine JP, Stamm WE, Suchland RJ, Rockey DD. 1998. Chlamydia trachomatis IncA is localized to the inclusion membrane and is recognized by antisera from infected humans and primates. Infect Immun 66:6017–21.

20. Bannantine JP, Griffiths RS, Viratyosin W, Brown WJ, Rockey DD. 2000. A secondary structure motif predictive of protein localization to the chlamydial inclusion membrane. Cell Microbiol 2:35–47.

21. Bui DC, Jorgenson LM, Ouellette SP, Rucks EA. 2021. Eukaryotic SNARE VAMP3 Dynamically Interacts with Multiple Chlamydial Inclusion Membrane Proteins. Infect Immun 89.

22. Stanhope R, Flora E, Bayne C, Derre I. 2017. IncV, a FFAT motif-containing Chlamydia protein, tethers the endoplasmic reticulum to the pathogen-containing vacuole. Proc Natl Acad Sci U S A 114:12039–12044.

23. Derre I, Swiss R, Agaisse H. 2011. The lipid transfer protein CERT interacts with the Chlamydia inclusion protein IncD and participates to ER-Chlamydia inclusion membrane contact sites. PLoS Pathog 7:e1002092.

24. Scidmore MA, Hackstadt T. 2001. Mammalian 14-3-3beta associates with the Chlamydia trachomatis inclusion membrane via its interaction with IncG. Mol Microbiol 39:1638–50.

25. Rzomp KA, Moorhead AR, Scidmore MA. 2006. The GTPase Rab4 interacts with Chlamydia trachomatis inclusion membrane protein CT229. Infect Immun 74:5362–73.

26. Weber MM, Lam JL, Dooley CA, Noriea NF, Hansen BT, Hoyt FH, Carmody AB, Sturdevant GL, Hackstadt T. 2017. Absence of Specific Chlamydia trachomatis Inclusion Membrane Proteins Triggers Premature Inclusion Membrane Lysis and Host Cell Death. Cell Reports 19:1406–1417.

27. Moore ER, Ouellette SP. 2014. Reconceptualizing the chlamydial inclusion as a pathogen-specified parasitic organelle: an expanded role for Inc proteins. 4.

28. Olson MG, Widner RE, Jorgenson LM, Lawrence A, Lagundzin D, Woods NT, Ouellette SP, Rucks EA. 2019. Proximity Labeling To Map Host-Pathogen Interactions at the Membrane of a Bacterium-Containing Vacuole in Chlamydia trachomatis-Infected Human Cells. Infect Immun 87.

29. Rucks EA, Olson MG, Jorgenson LM, Srinivasan RR, Ouellette SP. 2017. Development of a Proximity Labeling System to Map the Chlamydia trachomatis Inclusion Membrane. 7.

30. Gettemans J, Van Impe K, Delanote V, Hubert T, Vandekerckhove J, De Corte V. 2005. Nuclear actin-binding proteins as modulators of gene transcription. Traffic 6:847–57.

31. Lee Y-H, Campbell HD, Stallcup MR. 2004. Developmentally Essential Protein Flightless I Is a Nuclear Receptor Coactivator with Actin Binding Activity. Molecular and Cellular Biology 24:2103–2117.

32. Archer SK, Behm CA, Claudianos C, Campbell HD. 2004. The flightless I protein and the gelsolin family in nuclear hormone receptor-mediated signalling. Biochem Soc Trans 32:940–2.

33. Suriano AR, Sanford AN, Kim N, Oh M, Kennedy S, Henderson MJ, Dietzmann K, Sullivan KE. 2005. GCF2/LRRFIP1 Represses Tumor Necrosis Factor Alpha Expression. Molecular and Cellular Biology 25:9073–9081.

34. Rikiyama T, Curtis J, Oikawa M, Zimonjic DB, Popescu N, Murphy BA, Wilson MA, Johnson AC. 2003. GCF2: expression and molecular analysis of repression. Biochim Biophys Acta 1629:15–25.

35. Khachigian LM, Santiago FS, Rafty LA, Chan OL, Delbridge GJ, Bobik A, Collins T, Johnson AC. 1999. GC factor 2 represses platelet-derived growth factor A-chain gene transcription and is itself induced by arterial injury. Circ Res 84:1258–67.

36. Reed AL, Yamazaki H, Kaufman JD, Rubinstein Y, Murphy B, Johnson AC. 1998. Molecular cloning and characterization of a transcription regulator with homology to GC-binding factor. J Biol Chem 273:21594–602.

37. Broz P, Monack DM. 2013. Newly described pattern recognition receptors team up against intracellular pathogens. Nature Reviews Immunology 13:551–565.

38. Keating SE, Baran M, Bowie AG. 2011. Cytosolic DNA sensors regulating type I interferon induction. Trends in Immunology 32:574–581.

39. Yang P, An H, Liu X, Wen M, Zheng Y, Rui Y, Cao X. 2010. The cytosolic nucleic acid sensor LRRFIP1 mediates the production of type I interferon via a β-catenin-dependent pathway. Nature Immunology 11:487–494.

40. Bagashev A, Fitzgerald MC, Larosa DF, Rose PP, Cherry S, Johnson AC, Sullivan KE. 2010. Leucine-Rich Repeat (in Flightless I) Interacting Protein-1 Regulates a Rapid Type I Interferon Response. Journal of Interferon & Cytokine Research 30:843–852.

41. Wang T, Chuang TH, Ronni T, Gu S, Du YC, Cai H, Sun HQ, Yin HL, Chen X. 2006. Flightless I homolog negatively modulates the TLR pathway. J Immunol 176:1355–62.

42. Lee YH, Stallcup MR. 2006. Interplay of Fli-I and FLAP1 for regulation of beta-catenin dependent transcription. Nucleic Acids Res 34:5052–9.

43. Mills SJ, Ahangar P, Thomas HM, Hofma BR, Murray RZ, Cowin AJ. 2022. Flightless I Negatively Regulates Macrophage Surface TLR4, Delays Early Inflammation, and Impedes Wound Healing. Cells 11:2192.

44. Arora PD, Nakajima K, Nanda A, Plaha A, Wilde A, Sacks DB, McCulloch CA. 2020. Flightless anchors IQGAP1 and R-ras to mediate cell extension formation and matrix remodeling. Molecular Biology of the Cell 31:1595–1610.

45. Takimoto M. 2019. Multidisciplinary Roles of LRRFIP1/GCF2 in Human Biological Systems and Diseases. Cells 8:108.

46. Marei H, Carpy A, Woroniuk A, Vennin C, White G, Timpson P, Macek B, Malliri A. 2016. Differential Rac1 signalling by guanine nucleotide exchange factors implicates FLII in regulating Rac1-driven cell migration. Nat Commun 7:10664.

47. Mohammad I, Arora PD, Naghibzadeh Y, Wang Y, Li J, Mascarenhas W, Janmey PA, Dawson JF, McCulloch CA. 2012. Flightless I is a focal adhesion-associated actin-capping protein that regulates cell migration. FASEB J 26:3260–72.

48. Ariake K, Ohtsuka H, Motoi F, Douchi D, Oikawa M, Rikiyama T, Fukase K, Katayose Y, Egawa S, Unno M. 2012. GCF2/LRRFIP1 promotes colorectal cancer metastasis and liver invasion through integrin-dependent RhoA activation. Cancer Lett 325:99–107.

49. Kopecki Z, Cowin AJ. 2008. Flightless I: An actin-remodelling protein and an important negative regulator of wound repair. The International Journal of Biochemistry & Cell Biology 40:1415–1419.

50. Davy DA, Ball EE, Matthaei KI, Campbell HD, Crouch MF. 2000. The flightless I protein localizes to actin-based structures during embryonic development. Immunol Cell Biol 78:423–9.

51. Fukuhara S, Chikumi H, Gutkind JS. 2001. RGS-containing RhoGEFs: the missing link between transforming G proteins and Rho? Oncogene 20:1661–8.

52. Ouellette SP. 2018. Feasibility of a Conditional Knockout System for Chlamydia Based on CRISPR Interference. Front Cell Infect Microbiol 8:59.

53. Ouellette SP, Blay EA, Hatch ND, Fisher-Marvin LA. 2021. CRISPR Interference To Inducibly Repress Gene Expression in Chlamydia trachomatis. Infect Immun 89:e0010821.

54. Li GH, Arora PD, Chen Y, McCulloch CA, Liu P. 2012. Multifunctional roles of gelsolin in health and diseases. Med Res Rev 32:999–1025.

55. Liu YT, Yin HL. 1998. Identification of the binding partners for flightless I, A novel protein bridging the leucine-rich repeat and the gelsolin superfamilies. J Biol Chem 273:7920–7.

56. Strudwick XL, Cowin AJ. 2020. Multifunctional Roles of the Actin-Binding Protein Flightless I in Inflammation, Cancer and Wound Healing. Frontiers in Cell and Developmental Biology 8.

57. Fong KSK, De Couet HG. 1999. Novel Proteins Interacting with the Leucine-rich Repeat Domain of Human Flightless-I Identified by the Yeast Two-Hybrid System. 58:146–157.

58. Dai P, Jeong SY, Yu Y, Leng T, Wu W, Xie L, Chen X. 2009. Modulation of TLR signaling by multiple MyD88-interacting partners including leucine-rich repeat Fli-I-interacting proteins. J Immunol 182:3450–60.

59. Reuter J, Otten C, Jacquier N, Lee J, Mengin-Lecreulx D, Löckener I, Kluj R, Mayer C, Corona F, Dannenberg J, Aeby S, Bühl H, Greub G, Vollmer W, Ouellette SP, Schneider T, Henrichfreise B. 2023. An NlpC/P60 protein catalyzes a key step in peptidoglycan recycling at the intersection of energy recovery, cell division and immune evasion in the intracellular pathogen Chlamydia trachomatis. PLOS Pathogens 19:e1011047.

60. Mital J, Miller NJ, Fischer ER, Hackstadt T. 2010. Specific chlamydial inclusion membrane proteins associate with active Src family kinases in microdomains that interact with the host microtubule network. Cell Microbiol 12:1235–49.

61. Li Z, Chen C, Chen D, Wu Y, Zhong Y, Zhong G. 2008. Characterization of fifty putative inclusion membrane proteins encoded in the Chlamydia trachomatis genome. Infect Immun 76:2746–57.

62. Weber MM, Bauler LD, Lam J, Hackstadt T. 2015. Expression and localization of predicted inclusion membrane proteins in Chlamydia trachomatis. Infect Immun 83:4710–8.

63. Stephens RS, Kalman S, Lammel C, Fan J, Marathe R, Aravind L, Mitchell W, Olinger L, Tatusov RL, Zhao Q, Koonin EV, Davis RW. 1998. Genome sequence of an obligate intracellular pathogen of humans: Chlamydia trachomatis. Science 282:754–9.

64. Ende R, Derre I. 2019. A Coinfection Model to Evaluate Chlamydia Inc Protein Interactions. Methods Mol Biol 2042:205–218.

65. Han Y, Derre I. 2017. A Co-infection Model System and the Use of Chimeric Proteins to Study Chlamydia Inclusion Proteins Interaction. Front Cell Infect Microbiol 7:79.

66. Dehoux P, Flores R, Dauga C, Zhong G, Subtil A. 2011. Multi-genome identification and characterization of chlamydiae-specific type III secretion substrates: the Inc proteins. BMC Genomics 12:109.

67. Wang Y, Kahane S, Cutcliffe LT, Skilton RJ, Lambden PR, Clarke IN. 2011. Development of a transformation system for Chlamydia trachomatis: restoration of glycogen biosynthesis by acquisition of a plasmid shuttle vector. PLoS Pathog 7:e1002258.

68. Scidmore MA. 2005. Cultivation and Laboratory Maintenance of Chlamydia trachomatis. Curr Protoc Microbiol Chapter 11:Unit 11A 1.

69. Furness G, Graham DM, Reeve P. 1960. The Titration of Trachoma and Inclusion Blennorrhoea Viruses in Cell Cultures. Journal of General Microbiology 23:613–619.

70. Lucas AL, Ouellette SP, Kabeiseman EJ, Cichos KH, Rucks EA. 2015. The trans-Golgi SNARE syntaxin 10 is required for optimal development of Chlamydia trachomatis. Front Cell Infect Microbiol 5:68.

71. Wood NA, Blocker AM, Seleem MA, Conda-Sheridan M, Fisher DJ, Ouellette SP. 2020. The ClpX and ClpP2 Orthologs of Chlamydia trachomatis Perform Discrete and Essential Functions in Organism Growth and Development. mBio 11.

72. Ouellette SP, Gauliard E, Antosova Z, Ladant D. 2014. A Gateway((R)) – compatible bacterial adenylate cyclase-based two-hybrid system. Environ Microbiol Rep 6:259–67.

73. Karimova G, Pidoux J, Ullmann A, Ladant D. 1998. A bacterial two-hybrid system based on a reconstituted signal transduction pathway. Proc Natl Acad Sci U S A 95:5752–6.

